# Ketamine can produce oscillatory dynamics by engaging mechanisms dependent on the kinetics of NMDA receptors

**DOI:** 10.1101/2024.04.03.587998

**Authors:** Elie Adam, Marek Kowalski, Oluwaseun Akeju, Earl K. Miller, Emery N. Brown, Michelle M. McCarthy, Nancy Kopell

## Abstract

Ketamine is an NMDA-receptor antagonist that produces sedation, analgesia and dissociation at low doses and profound unconsciousness with antinociception at high doses. At high and low doses, ketamine can generate gamma oscillations (*>*25 Hz) in the electroencephalogram (EEG). The gamma oscillations are interrupted by slow-delta oscillations (0.1-4 Hz) at high doses. Ketamine’s primary molecular targets and its oscillatory dynamics have been characterized. However, how the actions of ketamine at the subcellular level give rise to the oscillatory dynamics observed at the network level remains unknown. By developing a biophysical model of cortical circuits, we demonstrate how NMDA-receptor antagonism by ketamine can produce the oscillatory dynamics observed in human EEG recordings and non-human primate local field potential recordings. We have discovered how impaired NMDA-receptor kinetics can cause disinhibition in neuronal circuits and how a disinhibited interaction between NMDA-receptor-mediated excitation and GABA-receptor-mediated inhibition can produce gamma oscillations at high and low doses, and slow-delta oscillations at high doses. Our work uncovers general mechanisms for generating oscillatory brain dynamics that differs from ones previously reported, and provides important insights into ketamine’s mechanisms of action as an anesthetic and as a therapy for treatment-resistant depression.

## Introduction

Ketamine is an N-methyl-D-aspartate receptor (NMDA_*R*_) antagonist that has analgesic, dissociative and hypnotic properties [1–4]. When administered at low doses, it causes analgesia and dissociation in patients and can be used for procedural sedation in a physician’s office or in the emergency room. When administered at high doses, it causes antinociception and profound unconsciousness and, therefore, can be used to create a state of general anesthesia in the operating room. In addition, ketamine is now being used as a therapy for treatment-resistant depression. Its antidepressant effects last long after the drug clears [5]. An excited brain state has been reported under these clinical conditions. At low doses, gamma oscillations (above 25Hz) can be expressed in the frontal electroencephalogram (EEG). At high doses, strong gamma oscillations are expressed in the frontal EEG and can be interrupted by slow-delta oscillations (below 4Hz) to produce down-states in the rhythmic activity [1].

Ketamine has been postulated to produce an excited brain state through disinhibition [6–8]. In particular, ketamine has been proposed to preferentially antagonize NMDA receptors of inhibitory neurons to drive a surge in excitation [6, 9–11]. It is in this disinhibited brain state that the oscillatory dynamics have been characterized [1, 3, 12]. However, taking into account this disinhibition, it remains unknown how the action of ketamine at the subcellular level can give rise to the oscillatory dynamics at the network level. We report neural circuit mechanisms that are engaged by NMDA_*R*_ antagonism under ketamine and that result in its oscillatory dynamics.

To examine these mechanisms, we developed a biophysical model of a cortical network exposed to ketamine. When we introduce ketamine as an NMDA_*R*_ antagonist, our model replicates the brain dynamics in human subjects and non-human primates administered ketamine and reveals brain mechanisms underlying the generation of these dynamics. Our model incorporates detailed kinetics for NMDA_*R*_ dynamics that underlie these mechanisms. By dissecting the model through simulations, we report three advances.

First, we have established that NMDA_*R*_ antagonism under ketamine can terminate spiking in active neurons with subthreshold background excitation. We find that, even when NMDA_*R*_s are antagonized non-preferentially across all neurons, global disinhibition can emerge when tonic inhibition is provided by interneurons with subthreshold background excitation or high resting membrane potential. Second, we have discovered an NMDA_*R*_-dependent mechanism that can generate gamma oscillations in a cortical network, through an interaction between NMDA- and GABA-receptor-mediated currents. As NMDA_*R*_ closing is known to be slow, this mechanisms stands in contrast to established mechanisms for gamma generation that rely on fast AMPA- and GABA-receptor transmission, such as the pyramidal-interneuronal gamma mechanism [13]. Third, we have discovered an NMDA_*R*_-dependent mechanism that can generate slow-delta oscillations in a cortical network, also through an interaction between NMDA- and GABA-receptor-mediated currents. This mechanism relies on an imbalance between excitatory and inhibitory activity, and stands in contrast to established mechanisms for slow-delta generation that rely on membrane channels, such as sodium-activated [14] and ATP-activated [15] potassium channels.

Through these findings, we explain how the effects of ketamine extend from the subcellular level to the network level to produce the oscillatory dynamics observed in brain activity. Our work proposes new general mechanisms for generating brain dynamics, provides new insight into the mechanism of action of ketamine, and suggests new mechanisms by which ketamine can provide therapeutic effects that go beyond anesthesia, dissociation and analgesia. Notably, ketamine has been established to produce antidepressant effect, that last long after drug clearance. Through our work, we find that the gamma oscillations could produce resonance in a subpopulation of interneurons expressing the vasoactive intestinal peptide. We believe that an excessive release of this peptide can trigger a cascade of synaptic changes and network reconfigurations that could enable or enhance the antidepressant effects of ketamine.

## Results

### Ketamine administration induces oscillatory dynamics in brain activity

In Figure 1A, a volunteer subject was administered a bolus of ketamine (SoI, red), with a dose high enough to produce general anesthesia (see **Materials and Methods**). The EEG of the subject shows a surge in gamma oscillations that coincides with a loss of response (LoR, yellow). These gamma oscillations appear in bursts and are interrupted by down-states in EEG activity. Throughout these bursts, the subject is unconsciousness. As the drug begins to clear from the system, brain dynamics typically transition from gamma oscillation bursts to stable gamma oscillations. These dynamics (Fig. 1A) are representative and are described in [1].

**Figure 1:**
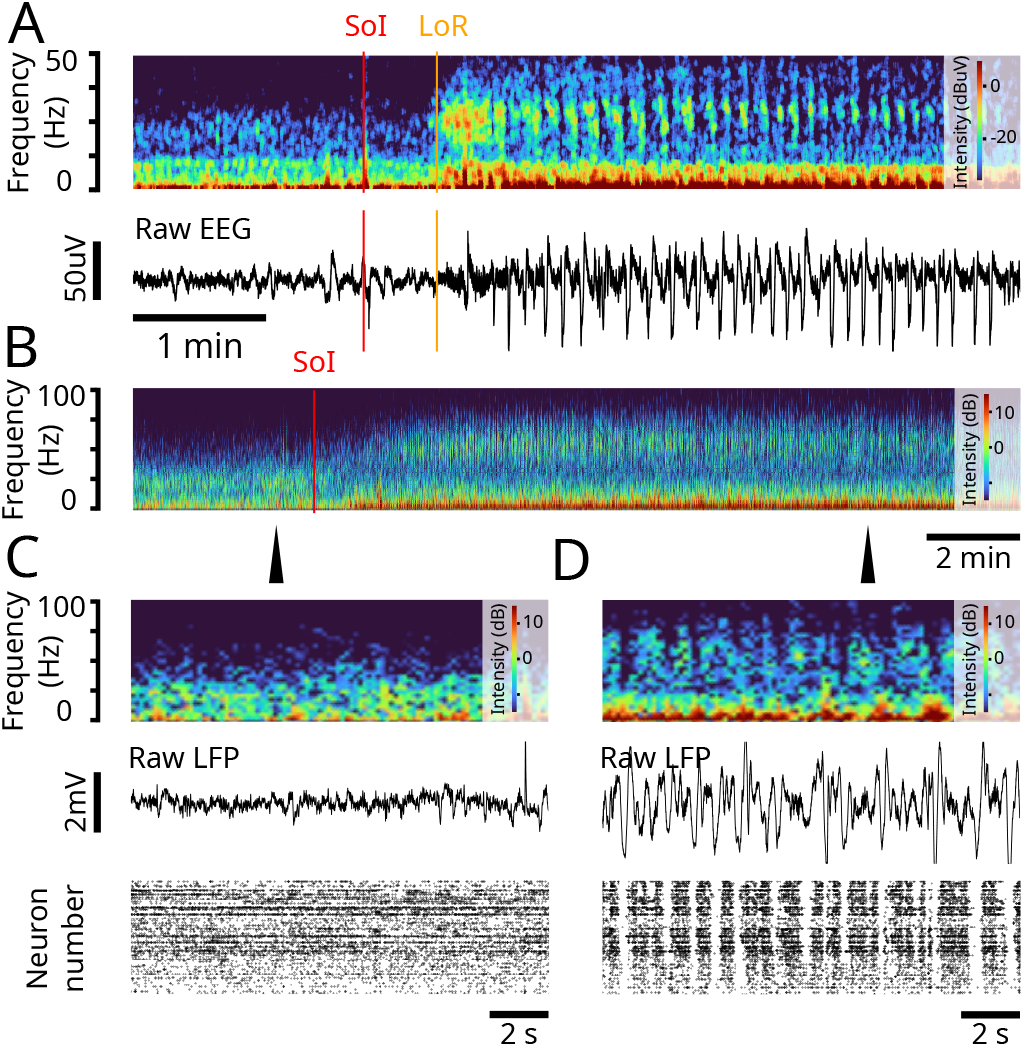
Ketamine produces gamma oscillations and up/down-states in humans and non-human primates. (Experimental data) **(A)** (top) Spectrogram of frontal EEG of a volunteer subject administered a bolus of ketamine. SoI (red) denotes the start of infusion. LoR (orange) denotes the loss of response. (bottom) Corresponding raw EEG. The red and orange lines denote the SoI and LoR, respectively. **(B)** Spectrogram of LFP recording from a nonhuman primate administered a ketamine bolus. SoI (red) denotes the start of infusion. **(C)** Close-up at the time-point indicated by the arrow in (B). (top) Raster plot of spiking activity before ketamine administration. (middle) LFP trace before before ketamine administration. (bottom) Spectrogram of LFP before ketamine administration. **(D)** Close-up at the time-point indicated by the arrow in (B). Same as (C) but after administering a bolus of ketamine.

Similar brain dynamics can be observed in non-human primates. In Figure 1B, a rhesus macaque was also administered a bolus of ketamine, with a dose high enough to produce general anesthesia (see **Materials and Methods**). The local field potential (LFP) in prefrontal cortex of the non-human primate shows a surge in gamma oscillations that become interrupted by down-states in LFP activity, where gamma oscillations are absent (Fig. 1B-D). The upand down-states observed in the LFP also extend to spiking activity. The down-states in gamma oscillations coincide with down-states in spiking activity (Fig. 1D.bottom). Throughout these upand down-states, the non-human primate is unconsciousness. These dynamics (Fig. 1B-D) are representative and are described in [16].

### Biophysical network modeling with detailed NMDA_*R*_ kinetics can produce the oscillatory dynamics under ketamine when NMDA_*R*_ are antagonized

To elucidate the brain mechanisms that generate the oscillatory dynamics under ketamine, shown in Figure 1 and observed more generally [1, 2], we developed a biophysical network model of a cortical circuit (Fig. 2A). The model consists of interacting excitatory pyramidal neurons (PYR) and inhibitory interneurons (IN-Phasic and IN-Tonic) (see **Materials and Methods**). While systemic administration of ketamine alters circuits throughout the brain, we focused on a minimal cortical network to show that elementary local network dynamics can give rise to the complexity of the dynamics under ketamine.

**Figure 2:**
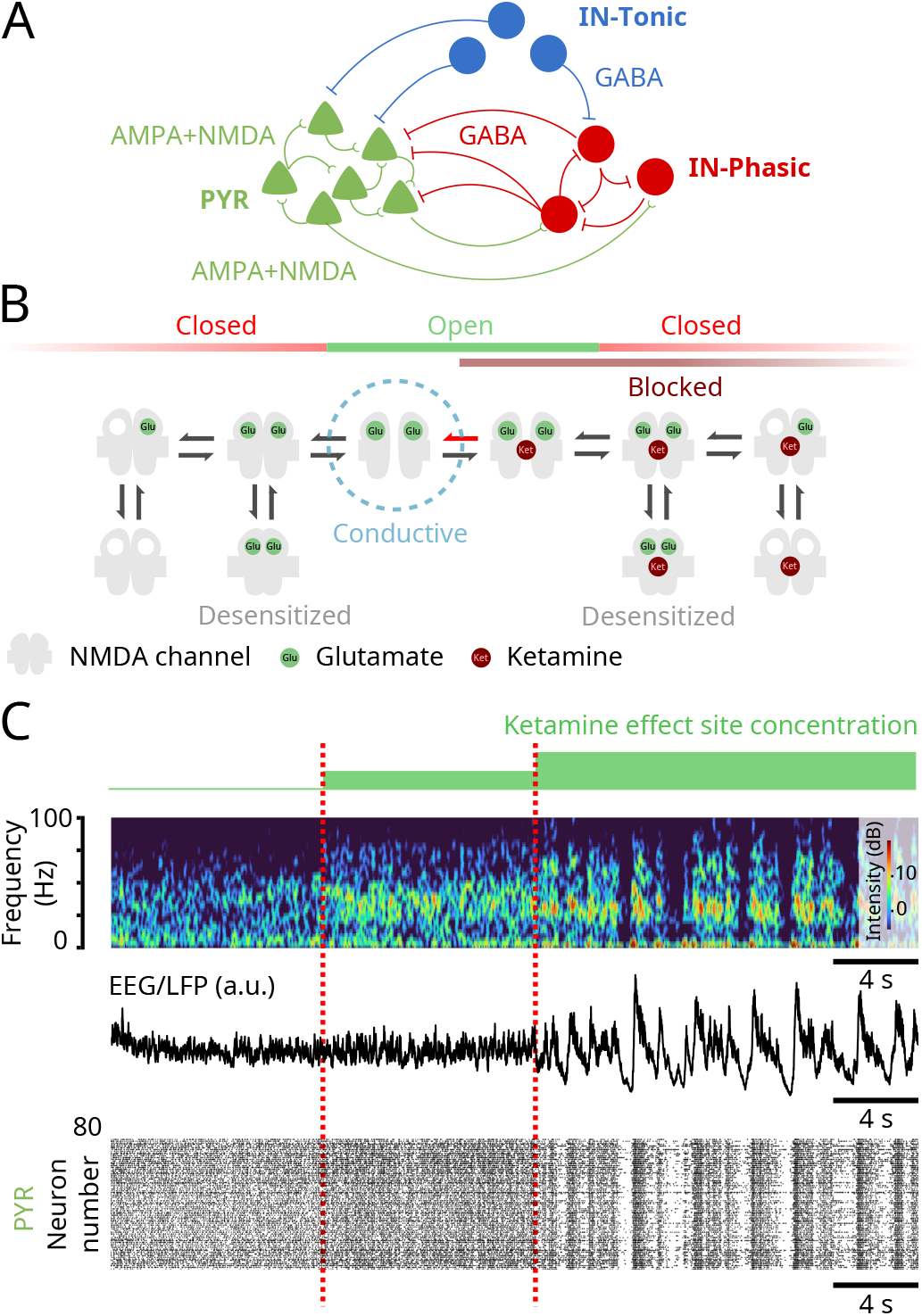
NMDA_*R*_ antagonism in a biophysical model reproduces the oscillatory dynamics under ketamine. (Model simulations) **(A)** Schematic of the biophysical network model. **(B)** Schematic of the 10-state model of NMDA_*R*_ kinetics. NMDA_*R*_ antagonism under ketamine was modeled as a decrease in the probability of NMDA_*R*_ channels unblocking (red arrow). **(C)** (top) Spectrogram of an EEG/LFP generated from a simulation of the biophysical model, under different effect site concentrations of ketamine. (middle) Corresponding EEG/LFP trace. (bottom) Corresponding raster plot of spiking activity.

Our findings are driven by a detailed model of NMDA_*R*_ kinetics and the biophysical effect of ketamine on these kinetics (Fig. 2B). We implemented a 10-state probabilistic model of NMDA_*R*_ kinetics, adapted from [17]. In this model, a closed NMDA_*R*_ channel can become open when two glutamate molecules bind to the receptor. However, an open channel does not always conduct ionic current. An open channel can be blocked with magnesium to become non-conductive. The channel becomes conductive once the magnesium is unblocked. Unblocking is a voltage-dependent mechanism and higher membrane potentials promote unblocking. NMDA_*R*_ channels can also close when blocked, a mechanism known as trapping. Finally, excessive usage of the receptor (channel blocked or unblocked) through glutamate binding can desensitize the receptor, and the channel enters a non-conducting state. This model then consists of 5 unblocked and 5 blocked states. More details on the NMDA_*R*_ kinetics are described in

## Materials and Methods

Ketamine, like magnesium, is an NMDA_*R*_ channel blocker. However, it is more effective at blocking than magnesium. To model different ketamine effect site concentrations, we decrease the rate of unblocking of the NMDA_*R*_ channel (Fig. 2B) (see **Materials and Methods**). As we increase ketamine effect site concentration, we find that the network exhibits the oscillatory dynamics characteristic of ketamine (Fig. 2C). At baseline, brain dynamics do not show preference to an oscillation. Introducing ketamine then gives rise to continuous gamma oscillations which, at higher effect site concentration, become interrupted by down-states in activity. These dynamics are observed at the level of the spectrogram (Fig. 2C.top), the simulated EEG/LFP (Fig. 2C.middle) and the spiking activity (Fig. 2C.bottom). These dynamics are preserved under randomized connectivity and initial conditions, as observed in 5 different simulations (Figs. S1-S5) in **Supplementary Information**.

While ketamine is primarily an NMDA_*R*_ antagonist, it is known to alter other channels such as hyperpolarization-activated nucleotide-gated (HCN) channels [18]. However, we decided to focus only on NMDA_*R*_ antagonism to show that, by itself, it can give rise to the range of brain dynamics under ketamine.

### Ketamine can decrease activity of interneurons by impairing the slow-unblock kinetics of NMDA_*R*_ channels to cause disinhibition

Despite antagonizing excitatory transmission, ketamine induces an excited brain state [2]. This phenomenon has been explained by the disinhibition hypothesis [6], in which ketamine is postulated to preferentially antagonize inhibitory neurons leading to greater excitation [7, 8]. To explain this antagonism, ketamine has been proposed to have a higher affinity to block subunits of NMDA_*R*_ channels preferentially expressed on inhibitory neurons [6, 9–11]. With our modeling, we find that even when NMDA_*R*_ kinetics are impaired equally among all neurons (without preferential targeting) some neurons with subthreshold background excitation can be shut down. When the neurons that are shut down are inhibitory, providing tonic inhibition onto circuits, this can lead to global disinhibition. As a first advance, we establish that NMDA_*R*_ antagonism can hyperpolarize neurons with subthreshold background excitation to stop them from firing, despite having their NMDA_*R*_ channels open, to be blocked or unblocked (Fig. 2B).

In our modeling, this shutdown depends on the mechanism of unblocking in NMDA_*R*_ channels (Fig. 3A). Unblocking is a voltage-dependent mechanism and admits two regimes: a fast unblock and a slow unblock [17]. High levels of depolarizations quickly relieve a magnesium block in NMDA_*R*_ channels. We call this a ‘fast unblock’. At low levels of depolarizations, such as subthreshold ranges, unblocking is still possible, although less likely. At these levels, unblocking of NMDA_*R*_ channels induces a small inward current that slowly depolarizes the neuron, that then leads to more unblocking and increases the inward current through positive feedback. We call this a ‘slow unblock’. By decreasing the probability of unblocking, ketamine impairs both fast and slow unblocks. In particular, ketamine decreases the inward current enabled by a subthreshold slow unblock. Neurons having subthreshold background excitations can rely on this current to sustain firing. In our model, these are represented by IN-Tonic neurons (see **Materials and Methods**). We find that by administering ketamine, we decrease the firing rate of INTonic neurons to a level at which we induce a complete shutdown (Fig. 3B). When tonic inhibition is mediated by such neurons, we find that ketamine leads to disinhibition.

**Figure 3:**
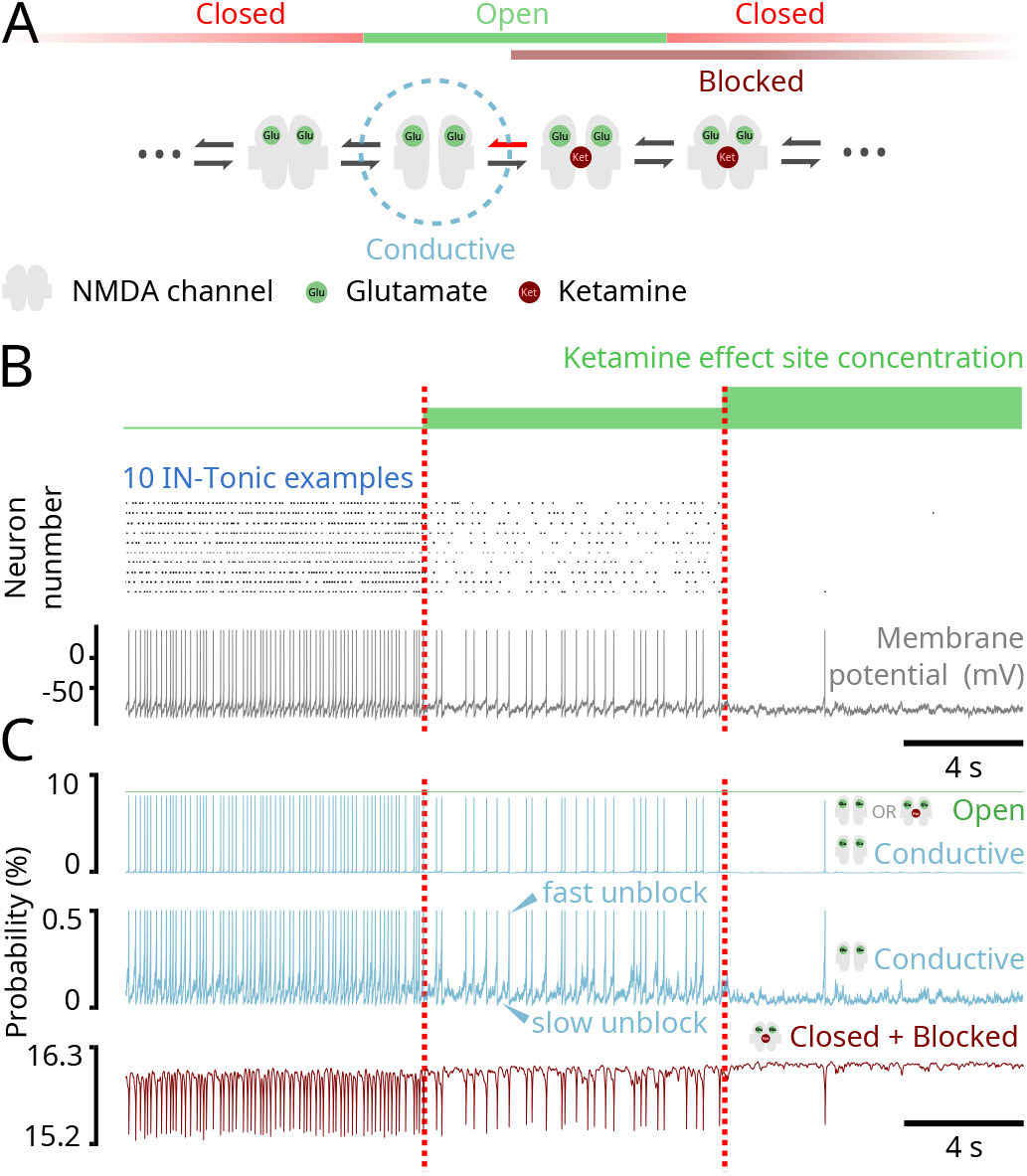
NMDA_*R*_ antagonism can shut down activity of neurons with subthreshold background excitation. (Model simulations) **(A)** Schematic showing parts of the 10-state model of NMDA_*R*_ kinetics. The red arrow represents the probability of unblocking, which was decreased. **(B)** (top) Raster plot of IN-Tonic activity at different ketamine effect site concentrations. Only 10 representative examples were selected. (bottom) Membrane potential of a representative example IN-Tonic neurons. **(C)** (top) Representative example from a neuron showing the probability of an NMDA_*R*_ channel being conductive (blue) and being open (blocked or unblocked; green) at different ketamine effect site concentrations. (middle) Scaled trace (blue) showing the probability of being conductive at different ketamine effect site concentrations. The slow ramp-up of probability indicates a slow-unblock (upper arrow) and a fast sudden-jump in probability indicates a fastunblock (lower arrow). (bottom) Representative example from the same neuron at top showing the probability of the NMDA_*R*_ channel being closed and blocked with 2 bound glutamate (brown).

The effect of ketamine on IN-Tonic neurons can be observed in the evolution of their NMDA_*R*_ channel state probabilities (Fig. 3C). The probability of an NMDA_*R*_ channel being conductive shows a stable fluctuation interrupted by sudden jumps that coincide with action potentials (Fig. 3C.top,middle). The stable fluctuations indicate periods of slow unblock during subthreshold membrane potentials, whereas the sudden surges indicate periods of fast unblock caused by high depolarization. The inward NMDA_*R*_ current is proportional to this probability (see **Materials and Methods**). As ketamine effect site concentration increases, we find that the probability of an NMDA_*R*_ channel being conductive decreases (Fig. 3C.top,middle), indicating a decrease in slow unblock current. However, this decrease occurs while the probabilty of a channel being open remains unchanged (Fig. 3C.top), indicating that the probability of being in a blocked state increases. Indeed, this decrease also coincides with an increase in the probability of NMDA_*R*_ channel being closed and blocked (Fig. 3C). Overall, this pushes neurons into a less excitable state.

We studied this effect of ketamine on network activity by providing IN-Tonic neurons with constant and maximal access to glutamate (see **Materials and Methods**). However, we also find that this effect is recreated when we modify the network to have IN-Tonic neurons directly receive local glutamate input from PYR neurons (Fig. S6A). Additionally, we find that the oscillatory dynamics (Fig. S6B.bottom) replicate those of the original model (Fig. 2A,B and Figs. S1-S5) as ketamine effect site concentration increases. Furthermore, our findings can also be replicated if subthreshold background excitation is replaced by high resting membrane potential (Fig. S6C,D) by altering the potassium leak current. Low-threshold spiking interneurons exhibit high resting membrane potentials [19] and include interneurons expressing somatostatin (SOM). In fact, SOM+ neurons have been found to be preferentially silenced by ketamine [20]. IN-Tonic neurons might comprise SOM+ neurons.

### Ketamine can engage an NMDA_*R*_-dependent network mechanism to generate gamma oscillations

A central signature of ketamine action is that the excitable brain state leads to an over-expression of gamma oscillations. Through our model, as a second advance, we find that NMDA_*R*_ antagonism by administering ketamine can yield gamma oscillations (Fig. 4A). As we will explain, we find that disinhibition allows NMDA_*R*_ channels to unblock at a gamma time-scale (Fig. 4B) and trigger bursts of spikes at a gamma time-scale in single neurons (Fig. 4C). These spikes are synchronized across neurons with GABA inhibition to form gamma oscillations (Fig. 4D). This mechanism is substantially different from pyramidal-interneuronal gamma (PING) that relies on the interaction of AMPAand GABA-receptormediated currents [13]. Indeed, we obtain the gamma oscillations in our model even in the absence of AMPA-receptors in the network (Fig. 4E).

**Figure 4:**
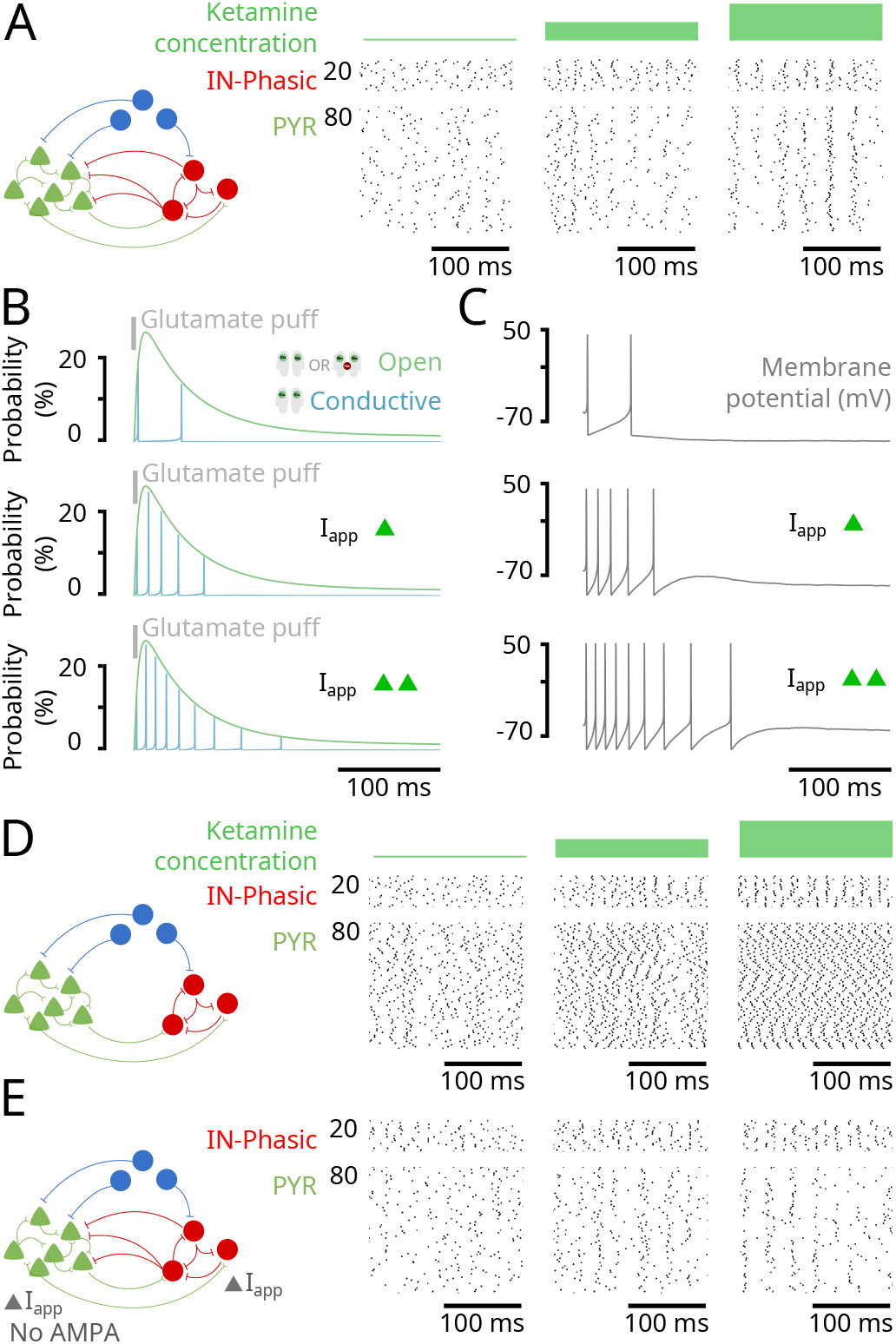
NMDA_*R*_ antagonism generates gamma oscillations through an NMDA_*R*_-dependent mechanism. (Model simulations) **(A)** (left) Schematic of the network in Figure 2. (right) Closed-up at the raster plot of spiking activity from the simulation in Figure 2 at different ketamine effect site concentrations. **(B)** Probability for an NMDA_*R*_ channel of an isolated neuron to be conductive (blue) and open (blocked or unblocked; green) following an initial puff of glutamate (grey), under different levels of background excitation (Iapp). **(C)** Membrane potentials (gray) corresponding to the conditions in (B). **(D)** (left) Schematic of the network where GABA input to PYR neurons was removed. (right) Raster plots of spiking activity for IN-Phasic and PYR neurons under the conditions in (left), at different ketamine effect site concentrations. **(E)** (left) Schematic of the network where AMPA receptors are removed from the network (while NMDA receptors are kept) and background current (Iapp) is adjusted to rectify the loss of excitation. (right) Raster plots of spiking activity for IN-Phasic and PYR neurons under the conditions in (left), at different ketamine effect site concentrations.

A core principle in this mechanism is that not all GABAergic neurons are inhibited under ketamine. In our model, some GABAergic neurons are disinhibited and then recruited to produce gamma oscillations. Indeed, once ketamine decreases the activity of IN-Tonic neurons, it relieves the inhibition onto PYR and IN-Phasic neurons. The slow unblock current of NMDA_*R*_ channels becomes more effective at depolarizing PYR and IN-Phasic neurons without this inhibition to counter it. Once an NMDA_*R*_ channel is open but blocked, depolarization can happen more rapidly, causing the membrane potential to reach the threshold and triggering an action potential. Specifically, by applying a puff of glutamate onto an isolated neuron (**Materials and Methods**), we find that, as we increased the background excitation of the neuron, NMDA_*R*_ channel unblocking is facilitated (Fig. 4B) and can cause a neuron to go from two spikes to a longer burst of spiking at a gamma time-scale (Fig. 4C). Through this mechanism triggered by disinhibition, NMDA_*R*_ channels can produce gamma oscillations at the level of a single neuron. The on-going excitatory activity in the network ensures that gamma bursts are expressed at the level of single neurons, PYR or IN-Phasic. But gamma oscillations at the level of a single neuron do not automatically translate to gamma oscillations at the level of the population. They need to be synchronized.

Synchrony is primarily achieved through inhibition from IN-Phasic neurons. Indeed, we find that if we remove the IN-Phasic to PYR projection, we lose synchrony in PYR neurons (Fig. 4D). However, by doing so, we keep the synchrony in IN-Phasic neurons (Fig. 4D). If we additionally remove IN-Phasic to IN-Phasic projections, we also lose synchrony in IN-Phasic neurons (Fig. S7A). Nevertheless, NMDA_*R*_ kinetics also have a role in ensuring synchrony. Once PYR neurons are synchronized, their activity also fosters synchrony in IN-Phasic neurons. Indeed, if we remove IN-Phasic to IN-Phasic projections while keeping IN-Phasic to PYR projections, we find that PYR synchrony is preserved and IN-Phasic synchrony is recovered (Fig. S7B). In fact, it is the synchronous opening of NMDA_*R*_ channels on INPhasic neurons that preserves the synchrony. Indeed, if we remove PYR input to IN-Phasic neurons and replace it by constant glutamate exposure, we lose the IN-Phasic synchrony (Fig. S7C). Whenever PYR neurons fire together, they re-open (blocked or unblocked) NMDA_*R*_ channels across all neurons, making it more likely for all neurons to immediately fire again, together.

### Ketamine can engage an NMDA_*R*_-dependent network mechanism to generate upand downstates

At high doses of ketamine, gamma oscillations are interrupted by down-states of activity. Through our model, as a third advance, we find that NMDA_*R*_ antagonism by administering ketamine can yield up- and down-states in activity (Fig. 5A). As we will explain, we find that it is the further impaired NMDA_*R*_ kinetics that push the dynamics into the slow-delta oscillatory regime.

**Figure 5:**
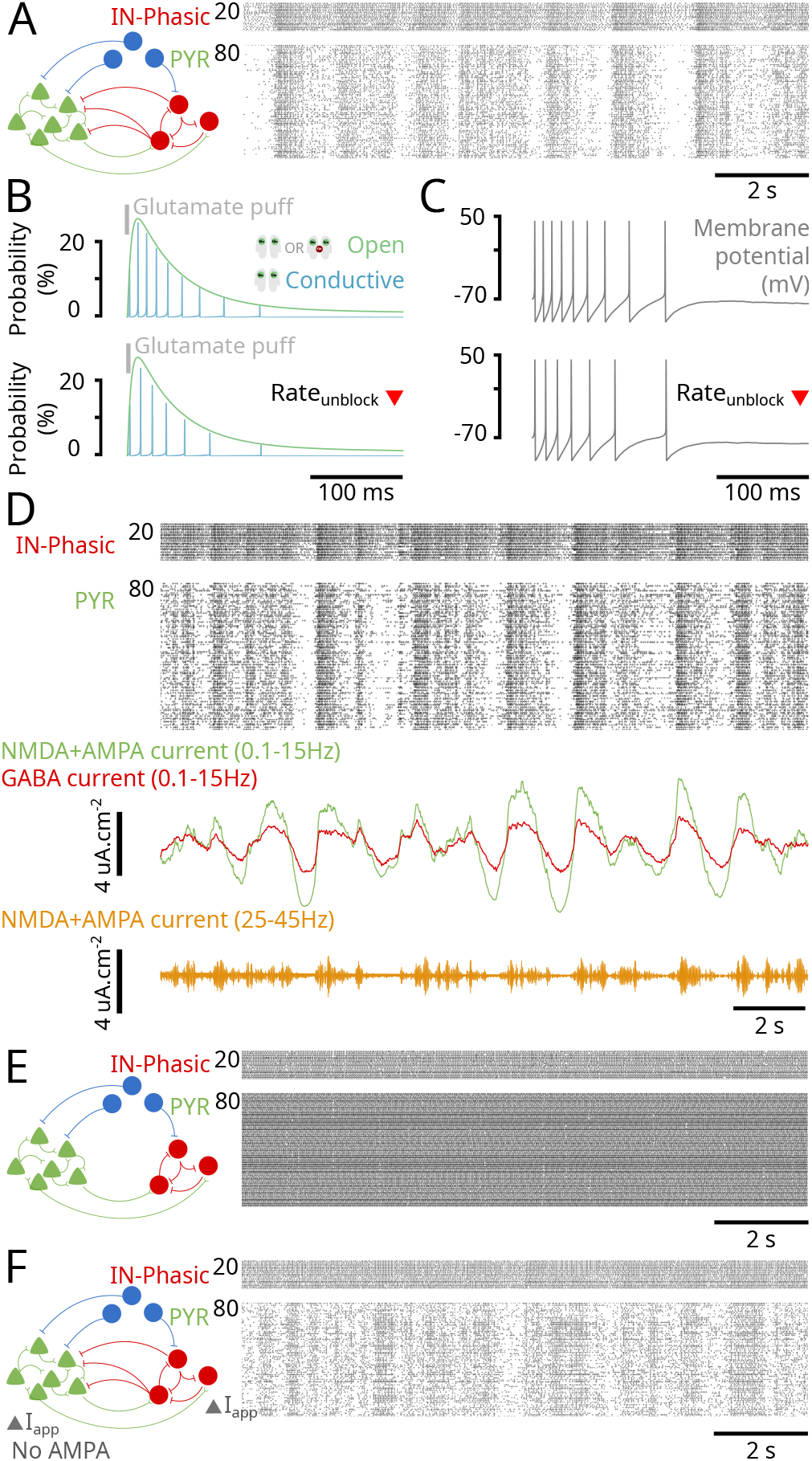
NMDA_*R*_ antagonism generates upand down-states through an NMDA_*R*_-dependent mechanism. (Model simulations) **(A)** (left) Schematic of the network in Figure 2. (right) Closeup at the raster plot of spiking activity for PYR and IN-Phasic neurons from the simulation in Figure 2, at high ketamine effect site concentration. **(B)** (top) Probability for an NMDA_*R*_ channel of an isolated neuron to be conductive (blue) and open (blocked or unblocked; green) following a puff of glutamate (grey), under high background excitation (disinhibited) at the baseline rate of unblocking for NMDA_*R*_ channels. (bottom) Same as (top) but using the high-dose rate of unblocking. **(C)** (top, bottom) Membrane potentials (gray) corresponding to the conditions in (B). **(D)** (top) Raster plot of spiking activity from Figure 2. (middle) Filtered excitatory (green) and inhibitory (red) currents input into PYR neurons. (bottom) Gamma oscillations in the EEG/LFP obtained through band-pass filtering. **(E)** (left) Schematic of the network where GABA input to PYR neurons was removed (right) Raster plots of spiking activity for IN-Phasic and PYR neurons under the conditions in (left), at high ketamine effect site concentration. **(F)** (left) Schematic of the network where AMPA receptors are removed (while NMDA receptors are kept) and background current (Iapp) is adjusted to rectify the loss of excitation. (right) Raster plots of spiking activity for IN-Phasic and PYR neurons under the conditions in (left), at high ketamine effect site concentration.

When ketamine is administered in our model, the probability of NMDA_*R*_ channels unblocking decreases. This has the effect of decreasing the slow unblock current of NMDA_*R*_s. For neurons with subthreshold background excitation that rely on it, this leads to a decrease in firing and eventual shutdown. For the remaining neurons, we find that this leads to a slowing of the timescale of the firing burst. Specifically, when unblocking decreases for isolated neurons, we find that the burst timescale slows down (Fig. 5B,C). In such a situation, the gamma oscillations in PYR neurons cannot be sustained indefinitely under gamma inhibition from IN-Phasic neurons.

This weakening produces up- and down-states following three steps. First, gamma oscillations are weakened and the activity of PYR neurons decreases with time, up to a point where it ceases. Second, once PYR activity decreases and ceases, IN-Phasic activity that is momentarily sustained by NMDA_*R*_ kinetics begins to decrease due to the absence of glutamate. Third, once IN-Phasic activity decreases enough, the inhibition onto PYR weakens, and we see re-emergence of PYR activity. These fluctuations are observed in the average excitatory (green) and inhibitory (red) currents in Figure 5D. In this cycle, inhibition from IN-Phasic neurons also plays a key role. Without it, synchrony is lost and the PYR neurons continuously spike (Fig. 5E). These up- and down-states also do not rely on AMPA receptors, as we can recreate them by removing AMPA receptors from the network (Fig. 5F). AMPA receptors do however play a role in enabling sharp transitions between the up- and down-states (e.g., compare Figure 5F to Figure 5A) due to their opening and closing kinetics that are faster than those of NMDA receptors.

### Ketamine can engage VIP+ neurons through resonance to gamma oscillations

Many interneurons that express the vasoactive intestinal peptide (VIP) are thought to express D-type potassium channels that conduct a D-current [21]. Through this D-current, we know that these VIP+ neurons can naturally produce bursts of gamma oscillations [22, 23]. This raises the possibility that the gamma oscillations produced by ketamine can recruit these VIP+ neurons through resonance. To investigate this, we modeled VIP+ neurons and introduced them into the network (Fig. 6A). When isolated VIP+ neurons receive a sinusoidal current with constant amplitude but increasing frequency (ZAP current), we find that they produce a burst of gamma in the range 30-40 Hz where gamma oscillations under ketamine are expressed (Fig. 6B). When we introduce VIP+ neurons in the network to receive PYR input, we find that they are entrained by PYR neurons (Fig. 6C). By examining membrane potentials, we find that VIP+ neurons spike sparsely at baseline and begin producing gamma bursts of oscillations as ketamine effect site concentration increases (Fig. 6C). Experimentally, the vasoactive intestinal peptide is found to have neurotrophic effects [24], and the rate of its release is tied to the stimulation frequency of VIP+ neurons [25]. Recruiting these neurons through resonance may underlie the long-term therapeutic effects of ketamine, particularly its anti-depressant effects.

**Figure 6:**
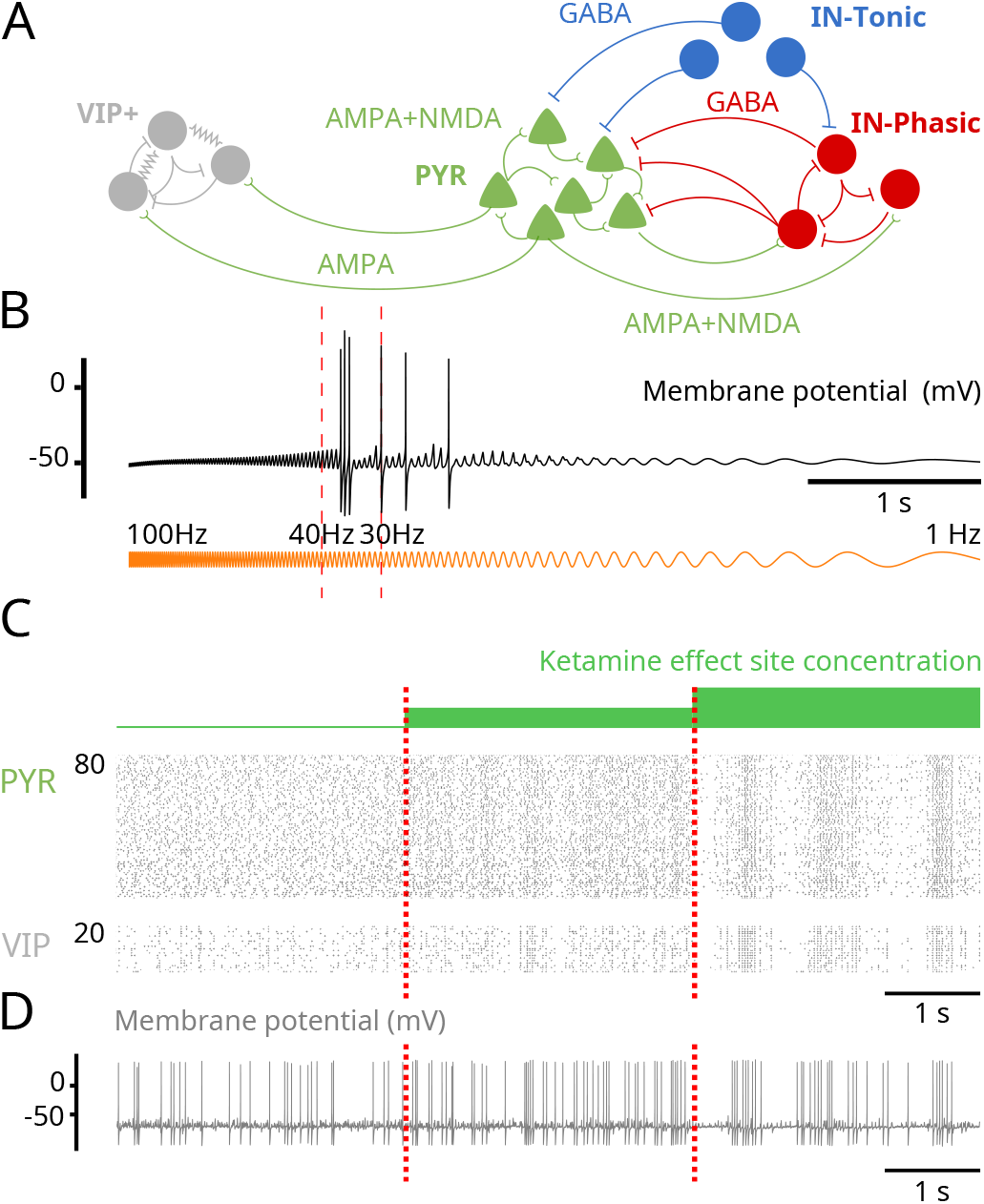
NMDA_*R*_ antagonism can engage VIP+ neurons through gamma resonance. (Model simulations) **(A)** Schematic of the augmented network including VIP+ neurons. **(B)** Membrane potential (black) of a representative VIP+ neuron in the network, where each receives a ZAP current (orange) as input instead of AMPA input from PYR neurons. The ZAP current sweeps from 100Hz to 1Hz and has constant amplitude throughtout. **(C)** Raster plots showing PYR and VIP+ neuron spiking activity at different effect site concentrations. **(D)** Membrane potential of a representative VIP+ neuron at different effect site concentrations.

## Discussion

### Ketamine can produce the oscillatory dynamics through NMDA_*R*_ antagonism

Using biophysical modeling, we have discovered mechanisms by which NMDA_*R*_ antagonism can generate the oscillatory brain dynamics observed under ketamine. These mechanisms connect the actions of ketamine at the subcellular level to the population activity through cellular and network effects. To derive them, we have focused on the action of NMDA_*R*_ antagonism on a cortical network. In our work, we report three advances.

First, we have established that NMDA_*R*_ antagonism can hyperpolarize neurons with subthreshold background excitation to terminate spiking under ketamine by impairing the ‘slow-unblock’ current of NMDA_*R*_ channels, despite these neurons being active before ketamine administration. We find that, even when NMDA_*R*_s are antagonized equally among all neurons, disinhibition can emerge when tonic inhibition is provided by interneurons with subthreshold background excitation or high resting membrane potential. The weakened activity of these interneurons can lead to global disinhibition.

Second, we have discovered an NMDA_*R*_-dependent mechanism that can generate gamma oscillations in a cortical network. Once the network is disinhibited by ketamine, we find that the gamma oscillations are generated at the single-neuron level by unblocking mechanisms in NMDA_*R*_ kinetics and are synchronized at the population level by GABA-receptor inhibition. This mechanism relies on the interaction of NMDA- and GABA-receptor-mediated currents and differs from known mechanisms such as the pyramidal-interneuronal gamma mechanism that relies on AMPA- and GABA-receptor-mediated currents [13].

Third, we have discovered an NMDA_*R*_-dependent mechanism that can generate slow-delta oscillations in a cortical network. Once NMDA_*R*_ kinetics are severely altered by high doses of ketamine, we find that up- and down-states emerge in three steps: pyramidal cell activity cannot be sustained and is shut down by inhibition, starting a down-state; interneuron activity which is sustained by NMDA_*R*_ currents begins to weaken after losing input from pyramidal cells; and once inhibition is weak enough, pyramidal cell activity re-emerges, starting an up-state. This mechanism also relies on the interaction of NMDA- and GABA-receptor-mediated currents and differs from known mechanisms reliant on membrane channels such as the sodium-activated [14] and ATP-activated [15] potassium channels.

Through these mechanisms, we find that the oscillatory dynamics could recruit additional populations of interneurons expressing VIP through gamma resonance. The increased activity of VIP+ neurons may increase the release of VIP. This peptide has neurotrophic effects [24] and its over-release may underlie the antidepressant effects of ketamine.

We believe that these mechanisms are more general and can be engaged in conditions beyond antagonized NMDA_*R*_. In fact, unblocking of NMDA_*R*_ is voltage-dependent and can then be modulated by changes in neuronal excitability. As a result, we expect a momentary disinhibition or activation of neuronal populations, which is possible in normal conditions, to engage these mechanisms to generate gamma or slow-delta oscillations. Therefore, our findings contribute an NMDA_*R*_ -centric viewpoint for generating brain oscillations in cortex that is complementary to what has been confined to AMPA- and GABA-receptor currents generating gamma oscillations and membrane channels generating slow-delta oscillations.

Our results rely on a detailed 10-state model of NMDA_*R*_ kinetics adapted from [17]. This level of detail allows us to directly impair the unblocking mechanism of NMDA_*R*_, by decreasing the rate of unblocking as a function of ketamine effect site concentration. This is in contrast to an alternate 4state models of NMDA_*R*_ that has been proposed but encapsulates the unblocking mechanism as a gating variable [26]. In fact, [27] examines mechanisms of ketamine action by decreasing the maximal conductance of NMDA_*R*_ channels, as modeled by [26], as a function of ketamine effect site concentration. This leaves NMDA_*R*_ kinetics unchanged under ketamine. We believe that NMDA_*R*_ channel unblocking and trapping mechanisms, and their effects on kinetics, are essential to ketamine action and should be explicitly considered.

### Therapeutic effects through gamma resonance under ketamine

Administering ketamine at low dose has been found to have antidepressant effects [5]. These effects are also long lasting. They have been proposed to emerge from a number of mechanisms: NMDA_*R*_ antagonism [5], activation of AMPA receptors [28], ketamine metabolites [29] and effects on opioid receptors [30]. Downstream of these mechanisms, the antidepressant effects have been found to need brain-derived neurotrophic factor (BDNF) signaling [31, 32]. Under ketamine, BDNF may be activated through a number of mechanisms to trigger the antidepressant effects.

Through our model, we find that the gamma oscillations could recruit neurons expressing the vasoactive intestinal peptide (VIP) through resonance to produce gamma bursts. This prediction would need to be verified experimentally. If correct, it would suggest a network mechanisms for ketamine’s antidepressant effect. We know that VIP+ neurons co-release VIP alongside GABA [33]. The rate of VIP release also increases with the rate of stimulation of VIP+ neurons [25]. These peptides have been found to have neurotrophic effects [24] that may lead to activation of BDNF [34]. The over-release of this peptide can trigger a cascade of synaptic changes and network reconfigurations that could enable or enhance the anti-depressant clinical effects under ketamine.

If this is true, then the gamma oscillations generated by ketamine would be critical for its antidepressant effects. Enhancing gamma oscillations under ketamine to further activate VIP+ neurons may then enhance ketamine’s therapeutic effects. Finally, that VIP+ activation is neurotrophic suggests that ketamine may be used as a therapy for conditions beyond treatment-resistant depression.

### Additional mechanisms for generating oscillatory dynamics under ketamine

In our work, we showed how NMDA_*R*_ antagonism in a simple cortical network can give rise to gamma and slow-delta oscillations. However, ketamine can have molecular effects other than NMDA_*R*_ antagonism. Furthermore, systemic administration of ketamine will affect all structures in the central nervous system.

The interaction between the cortex, the thalamus and the brainstem can contribute to slow-delta oscillatory dynamics. Indeed, removing significant excitatory input from the brainstem to the cortex can result in slow-delta oscillations [35]. Ketamine can also decrease the activity in excitatory arousal pathways from the parabrachial nucleus and the medial pontine reticular formation in the brainstem to the thalamus and basal forebrain through NMDA_*R*_ antagonism [36]. The coordination of down-states observed between cortex and thalamus suggests a mechanistic coordination between these two structures [16]. All of these indicate that other slow-delta oscillatory mechanisms can come into play to enhance or complement the mechanisms mediated by NMDA_*R*_ antagonism at the cortical level.

Ketamine can also alter hyperpolarization-activated nucleotide-gated (HCN) channels [18] for slow-delta oscillatory contributions. In fact, low doses of ketamine have been found to produce 3 Hz oscillations in posteromedial cortex [3, 12]. These 3 Hz oscillations have been found to rely on HCN channels [3]. Ketamine may act on these channels either indirectly through NMDA_*R*_ antagonism [3] or directly by inhibiting them [18] to produce these oscillations. Again, these mechanisms can enhance or complement the mechanisms mediated by NMDA_*R*_ antagonism at the cortical level.

### Disinhibition under ketamine

Although ketamine antagonizes excitatory transmission, it induces an excited brain state. To explain this, ketamine has been proposed to preferentially target inhibitory neurons, decreasing their activity and causing disinhibition [6–8]. To explain this targeting, ketamine has been proposed to have a higher affinity to block subunits of NMDA_*R*_ channel preferentially expressed on inhibitory neurons [6, 9–11]. However, through our modeling we find that disinhibition can also be realized without preferential targeting of NMDA_*R*_ on inhibitory neurons. Neurons, excitatory or inhibitory, with subthreshold background excitation that rely on their slow-unblock current to fire can stop firing under ketamine despite being heavily active before administration. These neurons are the IN-Tonic neurons in our model.

Background excitation for a neuron can be set by network dynamics and connections. As a result, different neuronal populations may be positioned to have different levels of background excitation through synaptic connections. Membrane channels also alter neuronal excitability and different populations of neurons can express different channels. For instance, SOM+ neurons are low-threshold spiking neurons known to have high resting membrane potentials [19]. They have also been found to be preferentially silenced by ketamine [20]. The IN-Tonic population in our network model may be comprised of SOM+ neurons.

While the disinhibition hypothesis suggests a weakening of GABAergic activity under ketamine, not all inhibitory neurons have to be weakened. Emerging experimental evidence shows that some inhibitory neurons are activated under ketamine [37]. In fact, our modeling suggests that some inhibitory neurons are activated and recruited to produce the gamma oscillations under ketamine. These neurons are the IN-Phasic neurons in our model.

### Altered balance of inhibition and excitation under ketamine

Overall, our mechanisms suggests that ketamine disrupts and alters the balance in excitation and inhibition as a function of dosage, producing switches in neuronal states. As the ketamine dose increases, the balance is first tilted to excess excitation then to more balanced inhibition at an excited regime to enable transitions between up- and down-states. The tilt in balance and switches in neuronal states set up the brain for the altered processing found under ketamine. The inhibition of neurons represented by IN-Tonic neurons can impair the brain’s ability to regulate dynamics. The disinhibition of the remaining population, represented by PYR or IN-Phasic neurons, can activate networks normally inhibited and enhance neurocognitive processes normally suppressed. The mechanisms that generate gamma oscillations also enhance synchronicity in neuronal populations. Spatially, this synchronicity can impair the formation of small natural cell assemblies and enhance the formation of large artificial ones. These changes in cell assemblies can alter perception and give rise to hallucinations [38–40]. Temporally, the synchronicity can override the expression of lower and higher frequency oscillations. This restricted expression of oscillations can stabilize dynamics, limiting the range of brain processing and leading to sedation. Finally, at high doses, the mechanisms that generates down-states in activity produce regular shutdowns across large populations of neurons. These down-states will regularly interrupt the continuity of brain processing and promote a state of unconsciousness. Ketamine’s dissociative effects have been tied to a 3 Hz oscillation in posteromedial cortex arising from altered HCN channels [3]. By altering the balance of inhibition and excitation, the mechanisms engaged by NMDA_*R*_ antagonism may interact with HCN channel alterations to further promote dissociation.

In future work, we will test experimentally how changes to the altered balance in excitation and inhibition under ketamine can change the expression of oscillations. Our findings suggest that gamma oscillations could be abolished by further shutting down inhibitory neurons, particularly neurons that represent IN-Phasic neurons. Our findings also suggest that slow-delta oscillations could also be abolished by causing further disinhibition in neuronal circuits. If this relationship between the effect of inhibition and the expression of oscillations is correct, then it can begin to inform us how other anesthetics, particularly GABAergic agents that potentiate GABAergic receptors, might interact with ketamine. Our findings also suggest a role for VIP+ neurons tied to the gamma oscillations. By recording from these neurons and manipulating them, we can begin to examine their effects on oscillatory dynamics and therapeutic effects. In summary, these next directions will examine approaches to alter the expression of oscillatory dynamics under ketamine which could change the effects of ketamine at a set dose for enhanced or different therapeutic effects.

## Material and Methods

### Human volunteer data

All data collection and experimental protocols in human subjects reported here were approved by the Mass General Brigham Human Research Committee (Institutional Review Board). All participants provided informed consent.

The EEG was acquired from a volunteer subject (26 year old male, 58.9 kg) administered solely ketamine to induce general anesthesia. Ketamine was administered as a single bolus of 2 mg/kg intravenously. The subject was instructed to click a mouse button when they heard auditory stimuli. Auditory stimuli were randomly presented every 4-8 seconds. All auditory stimuli were 1 second long and were delivered using headphones (ER2; Etymotic Research). Loss of response was determined once the subject stopped clicking the mouse button following the auditory stimuli. EEG was recorded using a 64-channel ANT-Neuro system (Ant-Neuro, Philadelphia, PA) sampled at 250 Hz. Channel F3 was selected and analyzed. The EEG signal was bandpass filtered between 0.1-50 Hz and plotted. The spectrogram was derived using the multitaper method [41].

### Non-human primate data

All procedures in non-human primates reported here followed the guidelines of the National Institutes of Health and were approved by the Massachusetts Institute of Technology’s Committee on Animal Care.

The local field potential (LFP) was recorded from a rhesus macaque (Macaca mulatta) aged 8 years (female, 6.6 kg). Ketamine was administered as a single 20 mg/kg bolus intramuscular dose. Fifteen minutes prior to ketamine administration, glycopyrrolate (0.01 mg/kg) was delivered to reduce salivation and airway secretions. The LFP was recorded from an 8 x 8 iridium-oxide contact microelectrode array (‘Utah array’, MultiPort: 1.0 mm shank length, 400 *µ*m spacing, Blackrock Microsystems, Salt Lake City, UT) implanted in the frontal cortex (vlPFC). The LFP was continuously recorded from 1–5 minutes prior to ketamine injection up to 18–20 minutes following ketamine injection. The LFP recorded at 30 kHz, was low-pass filtered to 250 Hz and then downsampled to 1 kHz. The LFP was bandpass filtered between 0.5-100 Hz using a 2nd order butterworth filter and plotted. The spectrogram was derived using the discrete fourier transform.

### Biophysical modeling

All neurons are modeled using a single compartment with Hodgkin-Huxley-type dynamics. The voltage change in each neuron is described by:

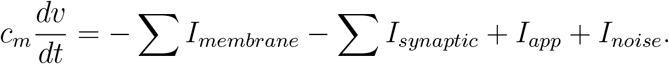

All neurons display a fast sodium current (*I*_*Na*_), a fast potassium current (*I*_*k*_), a leak current (*I*_*L*_) for membrane currents (*I*_*membrane*_). VIP+ neurons additionally displayed a D-current and an M-current. The synaptic currents (*I*_*synaptic*_) depend on the connectivity. The applied current (*I*_*app*_) is a constant that represents background excitation and the noise current (*I*_*noise*_) corresponds to a gaussian noise. The aggregate population activity (EEG/LFP) was defined as the sum of AMPA-receptor and NMDA-receptor synaptic currents into PYR neurons, bandpass filtered between 0.5-100 Hz. Modeling details and parameters are provided in **Supplementary Information**.

### Model of NMDA receptor kinetics

We implemented a 10-state probabilistic model of NMDA_*R*_ kinetics, adapted from [17]. The probabilty of being a certain state can be interpreted as the fraction of NMDA receptors in the particular state. State *i* transitions to state *j* with a rate *q*_*ij*_, denoting a conditional transition probability. The notation for the states and the rates are provided in Figure S8. If *Q* is the 10x10 transition matrix and *P* (*t*) is the probability vector of being in each of the 10 states, then:

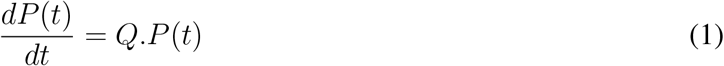

For example, for the conductive state O_AA_, we get:

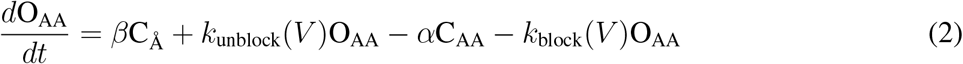

where:

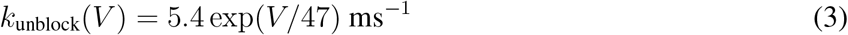

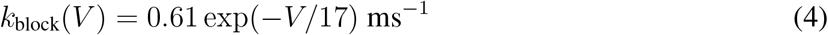

Each NMDA_*R*_ synaptic connection consists of such a probabilistic model. NMDA_*R*_ channels open following agonist (glutamate) binding. The concentration [Glu] denotes the amount of glutamate available at the synapse that can bind to NMDA receptors. Binding rates are determined by this concentration (Fig. S8). This concentration is given by:

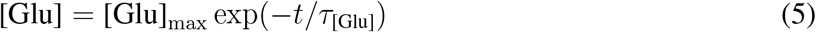

where *t* denotes the time of the last spike from the presynaptic neuron. We then modeled the NMDA current (*I*_*NMDA*_) as:

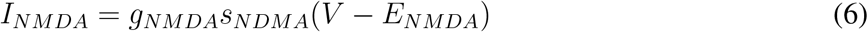

The gating variables *s*_*NDMA*_ is the sum:

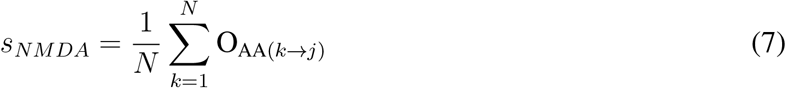

where *N* is the number of presynaptic neurons and 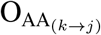 is the probability of being in state O_AA_ for NMDA_*R*_ synaptic connection *k → j*.

### Network connectivity

PYR and IN-Phasic neurons receive excitatory projections (AMPA- and NMDA-receptors) from PYR neurons. PYR and IN-Phasic neurons receive inhibitory projections (GABA-receptors) from IN-Phasic neurons. IN-Tonic neurons do not receive projections, but are modeled to have NMDA-receptors. The concentration [Glu] for NMDA_*R*_ is fixed constant for IN-Tonic neurons, to simulate constant input, which would allow NMDA_*R*_ kinetics to open and close. The concentration [Glu] for NMDA_*R*_ for PYR and IN-Phasic neurons is derived from pre-synaptic activity. We fixed the [Glu] concentration for IN-Tonic neurons to examine the slow-unblock current without closed-loop effects. However, the results are unchanged if that concentration is instead from pre-synaptic PYR activity (Fig. S6A).

### Modeling the effect of ketamine

The effect of ketamine was modeled by decreasing *k*_unblock_(0) from 5.4 ms^*−*1^ at baseline by 15% 4.6 ms^*−*1^ and then by 30% to 3.8 ms^*−*1^ at the highest effect site concentration. This decrease was applied to all NMDA receptors of all the neurons in the network.

### Modeling glutamate puffs on an isolated neuron

An isolated PYR neuron for NMDA_*R*_ kinetics simulations was formed by removing all projections. Glutamate puffs were simulated by setting [Glu] = 1 and letting the concentration decay following equation 5.

### Simulations and analysis

Our network model was programmed in C++ and compiled using GNU gcc. The differential equations were integrated using a fourth-order Runge-Kutta algorithm. The integration time step was 0.01ms. The model output was analyzed using Python 3.

## Acknowledgments

This work was generously supported by the JPB Foundation (E.N.B. and E.K.M.), the Picower Institute for Learning and Memory (E.N.B. and E.K.M.), the Simons Center for the Social Brain (E.K.M.) and the NIH Award P01 GM118269 (N.K., E.N.B. and E.K.M.).

## Author contributions

E.A., M.M., and N.K. designed the research; E.A. performed the research, with receptor and single-cell modeling contributions from M.K., and input from all authors; O.A. collected the human volunteer data; E.K.M. collected the non-human primate data; E.A., E.N.B., M.M. and N.K. wrote the manuscript with input from all authors.

## A Supplementary Information

### Biophysical modeling

All neurons were modeled using a single compartment with Hodgkin-Huxley-type dynamics. The voltage change in each neuron is described by:

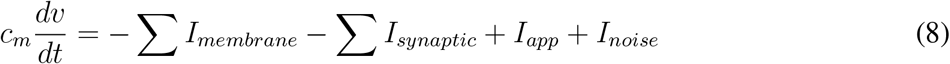

The membrane capacitance (*c*_*m*_) is normalized to 1 *µ*F *cm*^*−*2^ for all neurons. All neurons display a fast sodium current (*I*_*Na*_), a fast potassium current (*I*_*k*_), a leak current (*I*_*L*_) for membrane currents (*I*_*membrane*_). The synaptic currents (*I*_*synaptic*_) depend on the connectivity. The applied current (*I*_*app*_) is a constant that represents background excitation and the noise current (*I*_*noise*_) corresponds to a gaussian noise. The parameter values for PYR, IN-Phasic and IN-Tonic neurons generally follow the parameter values for excitatory and inhibitory neurons in [42] and [43]. They have been derived from experimental findings in the literature, as cited in previous work such as [13, 42, 43]. Any parameter value not within the ranges in [42] and [43] is justified when it is introduced, below.

### Membrane currents and background excitation

The basic membrane currents were modeled using Hodgkin-Huxley-type conductance dynamics and formulated as:

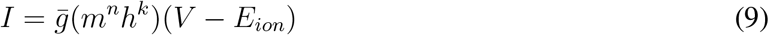

Every membrane current has a constant maximal conductance 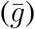 and a constant reversal potential (*E*_*ion*_). The activation (*m*) and inactivation (*h*) gating variables have *n*^*th*^ and *k*^*th*^ order kinetics with *n, k≥* 0. The dynamics of each gating variable evolves according to the kinetic equation (written here for the gating variable *m*):

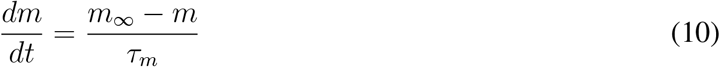

The steady-state function (*m*_*∞*_) and the time constant of decay (*τ*_*m*_) can be formulated as rate functions for each opening (*α*_*m*_) and closing (*β*_*m*_) of the ionic channel by using:

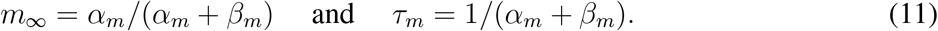

### Fast sodium current

The sodium current (*I*_*Na*_) has three activation gates (n=3) and only one inactivation gate (k=1). The rate functions for the sodium current activation (*m*) and inactivation (*h*) variables are formulated as:

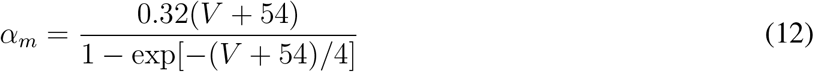

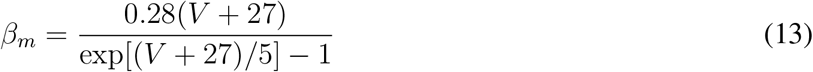

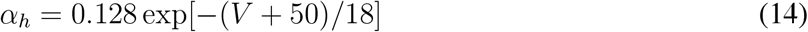

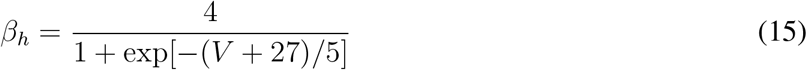

The maximal conductance of the sodium current is 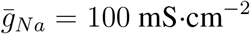. The sodium reversal potential is *E*_*Na*_ = 50mV.

### Fast potassium current

The fast potassium current (*I*_*K*_) has four activation gates (*n* = 4) and no inactivation gates (*k* = 0). The rate functions of the activation gate are described by:

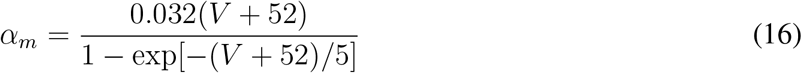

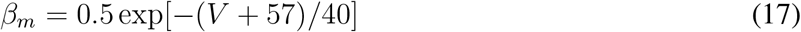

The maximal fast potassium channel conductance is 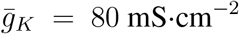. The reversal potential for potassium is *E*_*K*_ = *−*100mV.

### Leak current

The leak current (*I*_*L*_) has no gating variables (*n* = 0, *k* = 0). The maximal conductance of the leak channel is 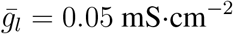. The leak channel reversal potential is *E*_*L*_ = *−*67mV.

### Applied current and noise

The baseline excitation and the sum of all excitatory and inhibitory exogenous inputs for a given neuron (e.g., from the cortex, thalamus and non-modeled input) is introduced into the model using a constant background excitation term (*I*_*app*_). To account for variability in background excitation, we further introduce a Gaussian noise term (*I*_*noise*_). The Gaussian noise has mean zero and standard deviation dependent on the neuronal cell type. The applied current (*I*_*app*_) is set to *−*0.25 *µ*A*·cm*^*−*2^ for PYR neurons, 0.1 *µ*A*·cm*^*−*2^ for IN-Phasic neurons, and *−*1.4 *µ*A*·cm*^*−*2^ for IN-Tonic neurons. The Gaussian noise (*I*_*noise*_) has mean 0 and standard deviation 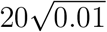 for PYR and IN-Phasic neurons and 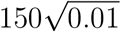 for IN-Tonic neurons where 0.01ms corresponds to the time step of integration in our simulations.

### Network structure and synaptic currents

Our model consisted of 80 PYR, 20 IN-Phasic, and 80 IN-Tonic neurons. Each PYR neuron receive 10 AMPA+NMDA projections from the remaining 79 PYR neurons, 10 GABA projections from the 20 IN-Phasic neurons and 5 GABA projections from the 80 IN-Tonic neurons. Each IN-Phasic neuron receive 10 AMPA+NMDA projections from 80 PYR neurons, 10 GABA projections from the remaining 19 IN-Phasic neurons and 5 GABA projections from the 80 IN-Tonic neurons. IN-Tonic neurons receive no projections in the base model. All projections were randomly and uniformly selected for a total of 10 from each cell type.

We needed IN-Tonic inhibition onto PYR and IN-Phasic neurons that is heterogenous and random, that would not foster synchrony and that would not pattern the activity of PYR and IN-Phasic neurons. To achieve that, we opted to provide GABAergic projections from a relatively large pool (80) of IN-Tonic neurons, instead of a small pool (e.g., 20) with size similar to that of IN-Phasic neurons. Each PYR and IN-Phasic neuron received input from 5 IN-Tonic neurons. This ensure that neurons (PYR and IN-Phasic) do not excessively have presynaptic input from the same combination of IN-Tonic neurons. As IN-Tonic neurons are not interconnected, the size of the IN-Tonic population only affects the heterogeneity and randomness of the inhibition onto PYR and IN-Phasic neurons.

While IN-Tonic neurons do not receive projections in the base model, they are modeled to have NMDA-receptors. The concentration [Glu] for NMDA_*R*_ of IN-Tonic neurons were all fixed to 1mM to provide constant excitatory input, which would allow NMDA_*R*_ kinetics to open and close. The concentration [Glu] for NMDA_*R*_ for PYR and IN-Phasic neurons is derived from pre-synaptic activity, as described in the next subsection. We fixed the [Glu] concentration for IN-Tonic neurons to examine the slow-unblock current without closed-loop effects. However, the results are unchanged if that concentration is instead from pre-synaptic PYR activity (Fig. S6A).

### NMDA-receptor kinetics and current

We implemented a 10-state probabilistic model of NMDA_*R*_ channel kinetics, adapted from [17]. The probabilty of being a certain state can be interpreted as the fraction of NMDA receptors in the particular state. State *i* transitions to state *j* with a rate *q*_*ij*_, denoting a conditional transition probability. The notation for the states and the rates are provided in Figure S8. If *Q* is the 10x10 transition matrix and *P* (*t*) is the probability vector of being in each of the 10 states, then:

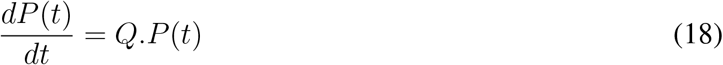

For example, for the conductive state O_AA_, we get:

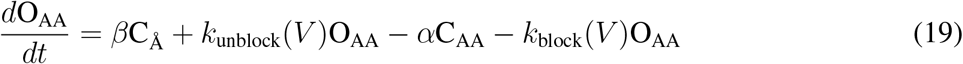

The values for the rates are:

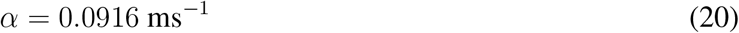

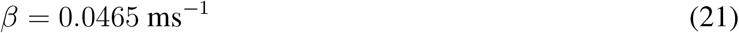

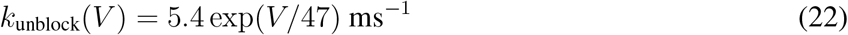

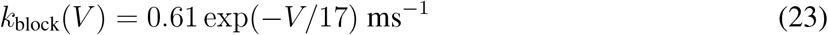

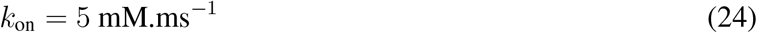

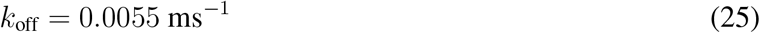

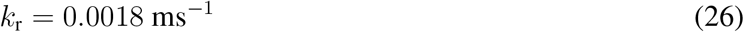

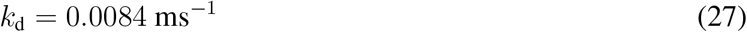

Built into the model of [17] is an asymmetry in the kinetics during the blocked versus unblocked regime. We decide to use a symmetric version of the model for simplicity and to have our results not depend on asymmetry. This distinction will not affect our results but, in general, can be a crucial component that can affect brain dynamics through NMDA-receptor kinetics. We also substituted the variable [Mg]^2+^ that appears in the blocking/unblocking rates of [17] by 1mM (see [17]).

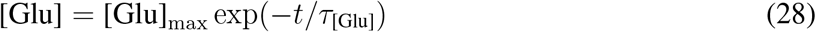

where *t* denotes the time of the last spike from the presynaptic neuron. We set [Glu]_max_ = 1mM and *τ*_[Glu]_ = 1.2ms. We then modeled the NMDA current (*I*_*NMDA*_) as:

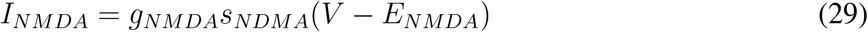

The gating variables *s*_*NDMA*_ is the sum:

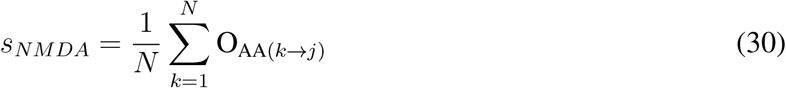

where *N* is the number of presynaptic neurons and 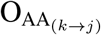 is the probability of being in state O_AA_ for NMDA_*R*_ synaptic connection *k → j*. The maximal conductance *g*_*NMDA*_ was set to be 8.5 mS.cm^*−*2^ for PYR neurons and 9.5 mS.cm^*−*2^ for IN-Phasic and IN-Tonic neurons. These values were chosen to preserve a reasonable NMDA-current/AMPA-current ratio as described in [44].

### GABA-and AMPA-receptor currents

We modeled GABA-currents (*I*_*GABAa*_) using a Hodgkin-Huxleytype conductance:

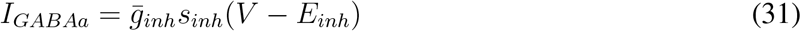

of the gating variables from all pre-synaptic connections. The gating variable *s*_*inh*_ for inhibitory GABAa synaptic transmission is the sum:

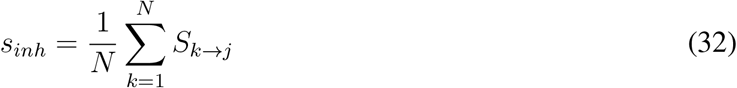

where *N* is the number of presynaptic neurons and *S*_*k→j*_ describes the kinetics of the gating variable, for each pair of presynaptic neuron *k* and postsynaptic neuron *j*, evolving according to:

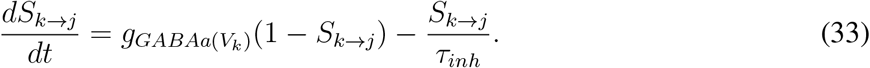

Note that *S*_*k→j*_ is a function of the presynaptic voltage *V*_*k*_, and its dynamics depend on the dynamics of the presynaptic neuron *k*. The rate functions for the open state of the GABAa receptor (*g*_*GABAa*_(*V*_*k*_)) is described by:

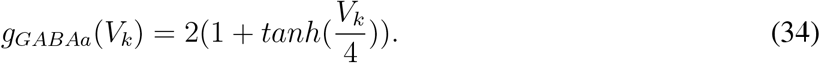

The maximal conductance 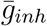 is scaled by the number of pre-synaptic neurons of the same type. Excitatory AMPA synaptic currents use the same set of equations as for the GABAa current with the “GABA” subscript replaced by “AMPA” and the “inh” subscript replaced by “exc”, with the exception for *g*_*AMPA*_(*V*_*k*_) modifed as 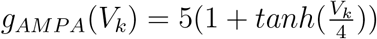.

We set *τ*_*inh*_ to be 6*ms* for projections from IN-Phasic neurons and 8*ms* for projections from IN-Tonic neurons. We set *τ*_*exc*_ = 1.5*ms* for all PYR projections. The potentials *E*_*inh*_ and *E*_*exc*_ were defined as *−*80*mV* and 0*mV*, respectively. Maximal conductances were 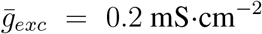 for AMPA-currents from PYR neurons, 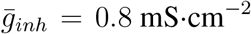 for GABA-currents from IN-Phasic neurons and 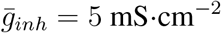 for GABA-currents from IN-Tonic neurons.

The value of the maximal conductance for IN-Tonic GABA-currents was selected so that IN-Tonic neurons can deliver adequate inhibition onto PYR and IN-Phasic neurons: abolishing gamma oscillation that emerge from PYR and IN-Phasic neuron disinhibition while maintaining spiking activity in PYR and IN-Phasic neurons. The value (5 mS .cm^*−*2^) appears to be much larger than that of IN-Phasic neurons (0.8 mS. cm^*−*2^). However, each PYR and IN-Phasic neuron received projections from 5 IN-Tonic neurons. These IN-Tonic neurons fire randomly, sparsely and not simultaneously. This is as opposed to IN-Phasic neurons which become highly active and synchronized when disinhibited. As a result, the maximal value for the conductance of IN-Tonic GABAergic currents during simulations is effectively near 1 mS .cm^*−*2^, the maximal conductance provided by each of the 5 IN-Tonic synaptic connections. Overall, the role of this IN-Tonic projection and its maximal conductance is to provide tonic inhibition onto PYR and IN-Phasic neurons to keep neuronal excitation low. These IN-Tonic GABAergic currents do not play a kinetic role in the generation of the gamma and slow-delta oscillations.

### Extended model with VIP neurons

We added 20 VIP neurons in an extended model. The properties and currents of these neurons can differ from what was already detailed above. The additions or changes are described below. The parameters for VIP neurons follow those presented for VIP in [45] and for fast spiking interneurons in [23].

### Modifications to membrane potentials

The fast sodium channel maximal conductance was set to 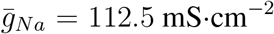 for VIP neurons. VIP neurons also displayed a D-current, an M-current and a modified fast potassium current.

### Fast potassium current for VIP+ neurons

The fast potassium current for VIP+ (*I*_*K*_) has two activation gates (n=2) and no inactivation gate (k=0). The steady state function for the potassium current activation (*n*) and its constant (*τ*_*n*_) are described by:

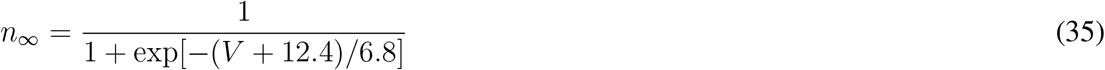

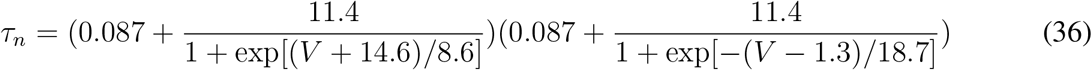

The maximal conductance of the potassium current is 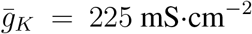. The potassium reversal potential is *E*_*Na*_ = *−*90mV. This instantiation of the current is only displayed in VIP neurons.

### D-current

The fast-activating, slowly inactivating potassium D-current (*I*_*D*_) is described mathematically as in [22] and has three activation gates (n = 3) and one inactivation (k = 1) gate. The steady state functions for the activation (m) and inactivation (h) variables and their time constants (*τ*_*m*_ and *τ*_*h*_, respectively) are described by:

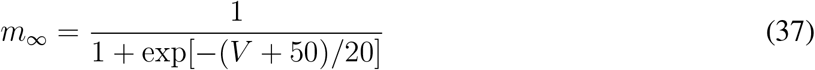

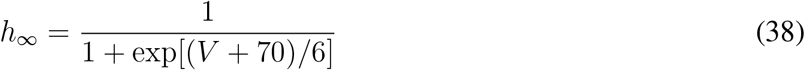

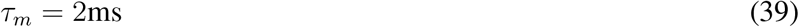

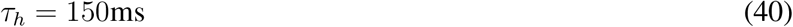

The maximal conductance of the D-current is 5.6 mS.cm^*−*2^ for VIP.

### M-current

The M-current (*I*_*M*_) has one activation gate (*n* = 1) and no inactivation gate (*k* = 0). The rate functions for the M-current activation gate are described by:

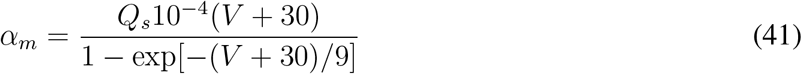

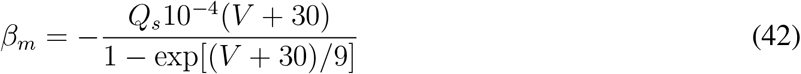

We use a *Q*_10_ factor of 2.3 to scale the rate functions of the M-current since the original formulation of these kinetics described dynamics at 23^*°*^*C* [46]. Thus, for a normal body temperature of 37^*°*^*C*, the M-current rate equations are scaled by *Q*_*s*_, which is formulated as:

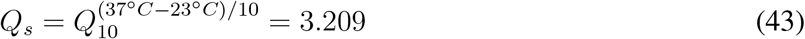

The maximal M-current conductance is 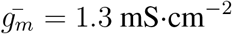 for VIP.

### Leak current

The maximal conductance of the leak channel is 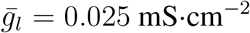 for VIP. The leak channel reversal potential is *E*_*L*_ = *−*70mV for VIP.

### Applied current and noise

The applied current (*I*_*app*_) is set to 5 *µ*A*·cm*^*−*2^ for VIP. The Gaussian noise (*I*_*noise*_) has mean 0 and standard deviation 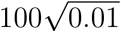 for VIP where 0.01ms corresponds to the time step of integration in our simulations.

### Modifications to connectivity

Maximal conductances was 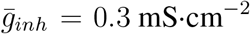 for GABA-currents from VIP projections and 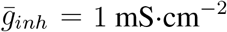 for AMPA-currents from PYR projections.

### Gap junctions

VIP neurons were additionally connected via electrical gap junctions. The electrical coupling of VIP neuron *j* to neuron *k* was defined as:

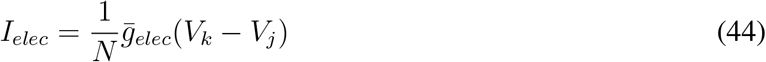

with *N* equal to the number of VIP neurons that *j* is coupled to. These coupling introduce a current ∑^*k*^ *I*_*elec*_ in VIP neuron *j*. We set 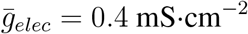

### Network and connectivity

Each VIP neuron receive 10 AMPA projections from PYR neurons and 10 GABA projections from VIP+ neurons. Each VIP+ neuron is also coupled to 10 remaining VIP neurons. All the projections are randomized. The projections received by PYR and IN-Phasic neurons are uniformly selected for a total of 10 from each cell type. The projections received by VIP+ neurons are selected by a bernouilli coin flip of probability 0.125 for AMPA projections from PYR neurons, 0.5 for GABA projections from VIP neurons, 0.5 for electrical coupling from VIP neurons. The electrical coupling of VIP+ neurons was made symmetrical.

### Modeling the effect of ketamine

The effect of ketamine was modeled by decreasing *k*_unblock_(0) from 5.4 ms^*−*1^ at baseline by 15% 4.6 ms^*−*1^ and then by 30% to 3.8 ms^*−*1^ at the highest dose. This decrease was applied to all NMDA receptors of all the neurons in the network.

### Network perturbation simulations

An isolated PYR neuron for NMDA_*R*_ kinetics simulations was formed by removing all projections, and setting [Glu] = 1 and letting it decay per the equation to simulate a glutamate puff. The applied current *I*_*app*_ was set to *−*2 *µ*A *·cm*^*−*2^, *−*1.25 *µ*A *·cm*^*−*2^ and *−*0.5 *µ*A*· cm*^*−*2^ in Figure 4B,C (top to bottom). The applied current *I*_*app*_ was set to *−*0.5 *µ*A *·cm*^*−*2^ and *k*_unblock_(0) decreased from 5.4ms to 3.8ms in Figure 5B,C (top to bottom).

In Figures 4F and 5G, *I*_*app*_ increased to 0.15 *µ*A*·cm*^*−*2^ for PYR neurons and to 0.6 *µ*A*·cm*^*−*2^ for IN-Phasic neurons. In Figure S6C, *I*_*app*_ was decreased to *−*2.05 *µ*A*·cm*^*−*2^ and *E*_*l*_ was increased to *−*53*mV* . In Figure S7B, *I*_*app*_ was decreased to *−*1.4 *µ*A*·cm*^*−*2^. In Figure S7C [Glu]was fixed to 0.5mM, and *I*_*app*_ was decreased to *−*0.9 *µ*A*·cm*^*−*2^.

### Aggregate population activity

Synaptic currents have been used in models of LFP and EEG [47]. We model the population aggregate activity (EEG/LFP) as the sum of all AMPA and NMDA currents going into PYR neurons. Thus our aggregate signal is tracking the excitatory activity driving spiking throughout the network.

### Simulation and analysis

Our network models were programmed in C++ and compiled using GNU gcc. The differential equations were integrated using a fourth-order Runge Kutta algorithm. The integration time step was 0.01 ms. Model output is graphed and analyzed using Python 3. Signals were filtered using a butterworth bandpass filter of order 2.

## B Supplementary Figures

**Figure S1:**
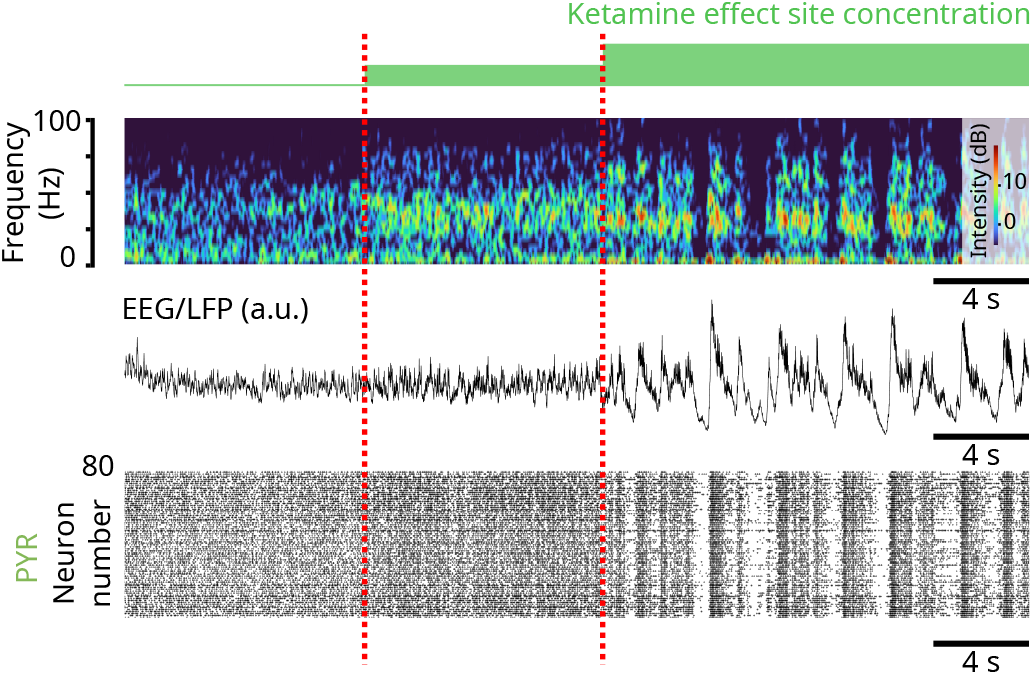
Simulation 1: NMDA_*R*_ antagonism in a biophysical model reproduces the oscillatory dynamics under ketamine. (Model simulations) (top) Spectrogram of an EEG/LFP generated from a simulation of the biophysical model, under different effect site concentrations of ketamine. (middle) Corresponding EEG/LFP trace. (bottom) Corresponding raster plot of spiking activity.

**Figure S2:**
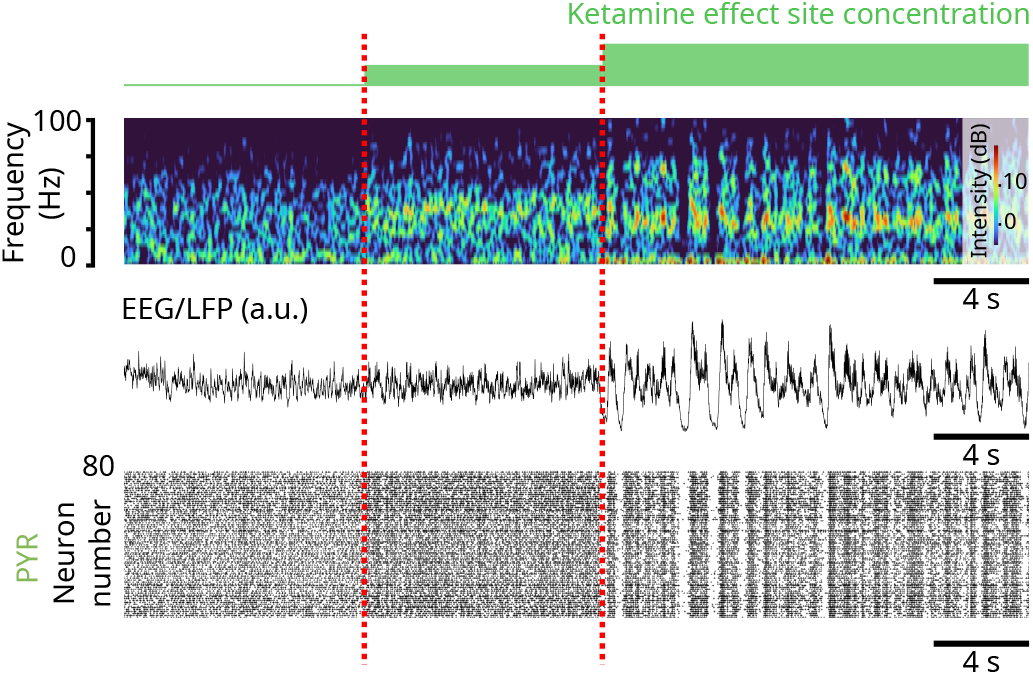
Simulation 2: NMDA_*R*_ antagonism in a biophysical model reproduces the oscillatory dynamics under ketamine. (Model simulations) (top) Spectrogram of an EEG/LFP generated from a simulation of the biophysical model, under different effect site concentrations of ketamine. (middle) Corresponding EEG/LFP trace. (bottom) Corresponding raster plot of spiking activity.

**Figure S3:**
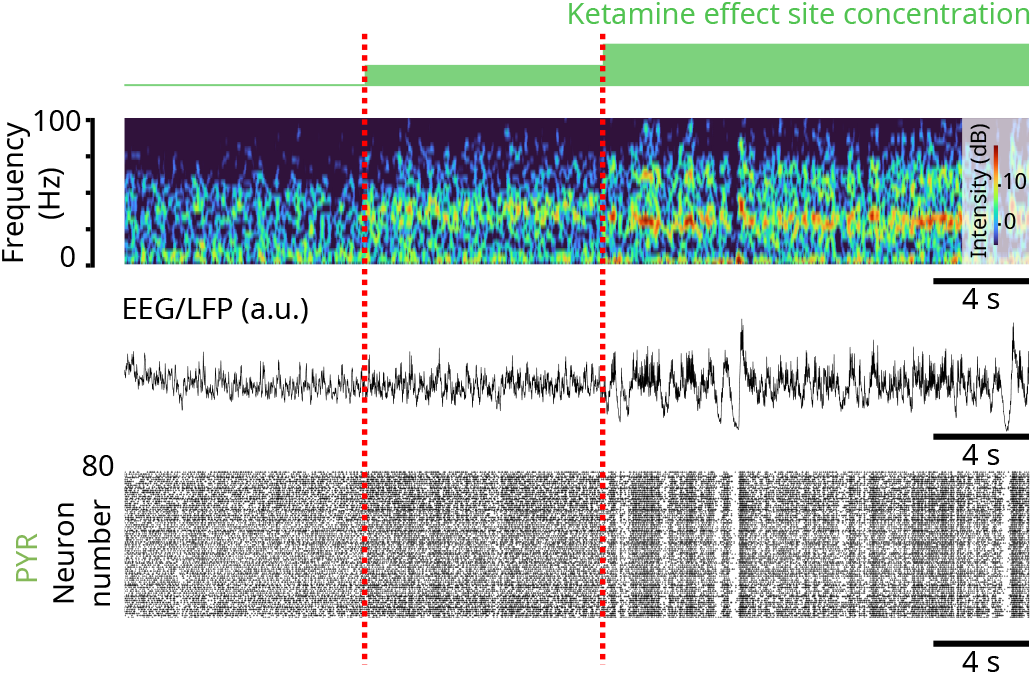
Simulation 3: NMDA_*R*_ antagonism in a biophysical model reproduces the oscillatory dynamics under ketamine. (Model simulations) (top) Spectrogram of an EEG/LFP generated from a simulation of the biophysical model, under different effect site concentrations of ketamine. (middle) Corresponding EEG/LFP trace. (bottom) Corresponding raster plot of spiking activity.

**Figure S4:**
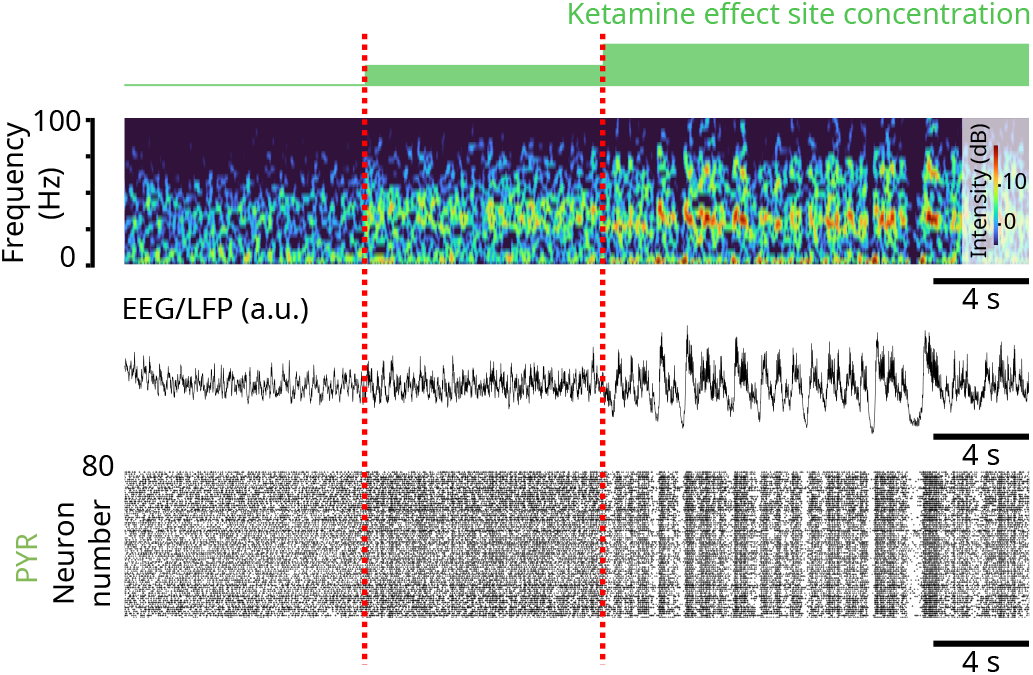
Simulation 4: NMDA_*R*_ antagonism in a biophysical model reproduces the oscillatory dynamics under ketamine. (Model simulations) (top) Spectrogram of an EEG/LFP generated from a simulation of the biophysical model, under different effect site concentrations of ketamine. (middle) Corresponding EEG/LFP trace. (bottom) Corresponding raster plot of spiking activity.

**Figure S5:**
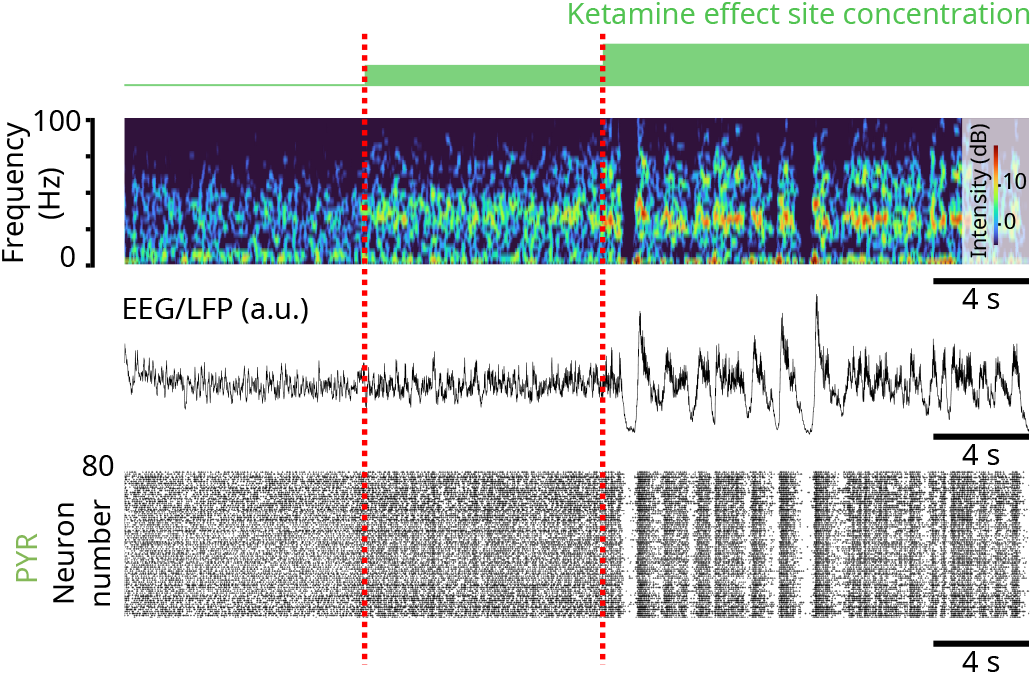
Simulation 5: NMDA_*R*_ antagonism in a biophysical model reproduces the oscillatory dynamics under ketamine. (Model simulations) (top) Spectrogram of an EEG/LFP generated from a simulation of the biophysical model, under different effect site concentrations of ketamine. (middle) Corresponding EEG/LFP trace. (bottom) Corresponding raster plot of spiking activity.

**Figure S6:**
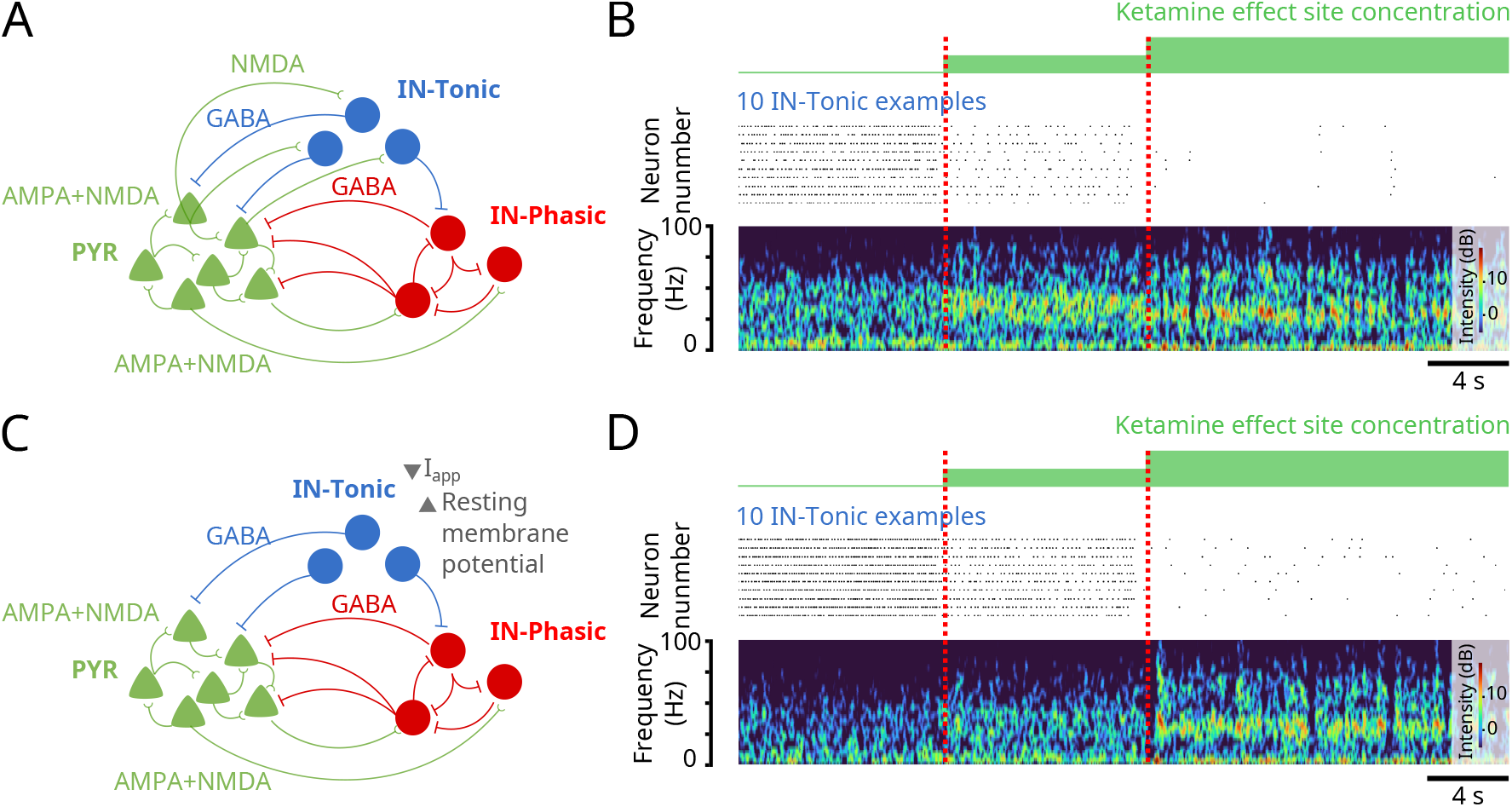
NMDA_*R*_ antagonism can shut down inhibition under different conditions. (Model simulations) **(A)** Schematic of the biophysical model with added NMDA projections from PYR to IN-Tonic neurons. **(B)** (top) Raster plot of spiking activity of representative IN-Tonic neurons. (bottom) Spectrogram of an LFP generated from a simulation of the biophysical model of (A), under different effect site concentrations of ketamine. **(C)** Schematic of the biophysical model where the background excitation is decreased and the resting membrane potential is increased. **(D)** (top) Raster plot of spiking activity of representative IN-Tonic neurons. (bottom) Spectrogram of an LFP generated from a simulation of the biophysical model of (C), under different effect site concentrations of ketamine.

**Figure S7:**
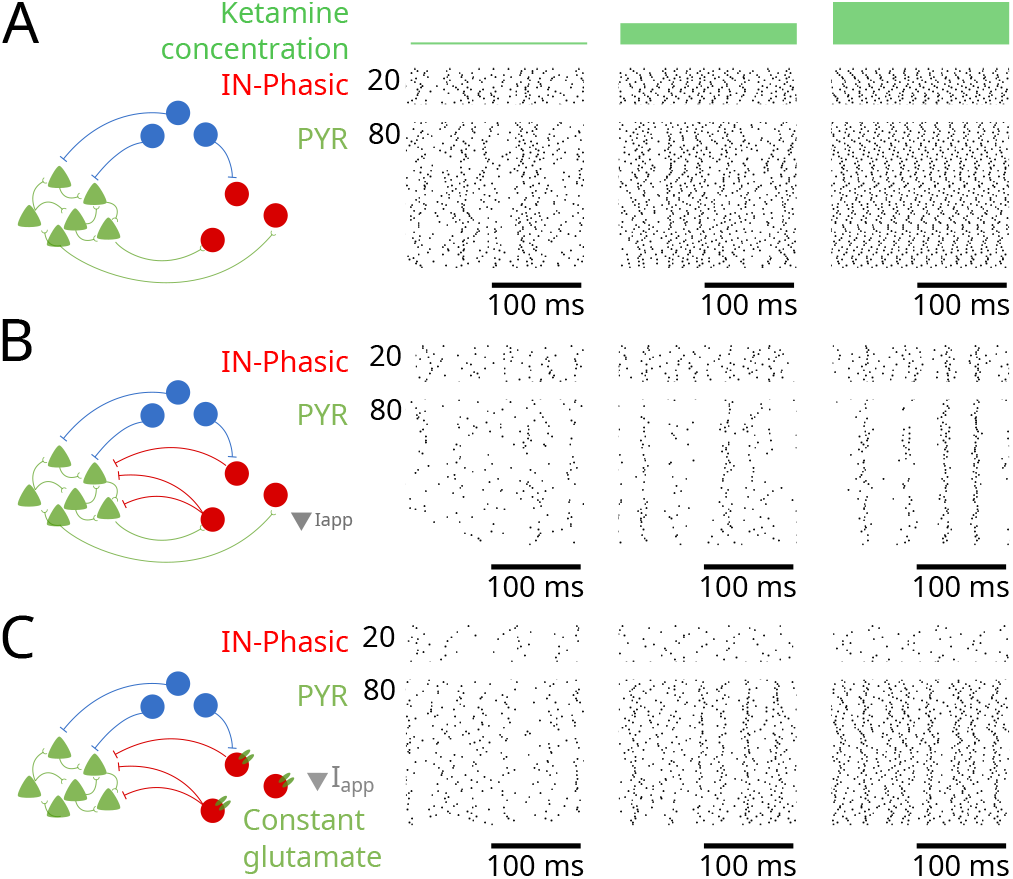
Effects of GABA-receptor inhibition on gamma oscillations under NMDA_*R*_ antagonism. (Model simulations) **(A)** (left) Schematic of the network for the simulation where IN-PYR and IN-IN projections were removed. (right) Raster plots of spiking activity for IN-Phasic and PYR neurons under the conditions in (left), at different ketamine effect site concentrations. **(B)** (left) Schematic of the network for the simulation where IN-IN projections were removed and Iapp adjusted accordingly to correct increase in excitation. (right) Raster plots of spiking activity for IN-Phasic and PYR neurons under the conditions in (left), at different ketamine effect site concentrations. **(C)** (left) Schematic of the network for the simulation where IN-IN and PYR-IN projections were removed. NMDA_*R*_ on IN were exposed to constant glutamate concentration, and Iapp was decreased to correct for increase of excitation. (right) Raster plots of spiking activity for IN-Phasic and PYR neurons under the conditions in (left), at different ketamine effect site concentrations.

**Figure S8:**
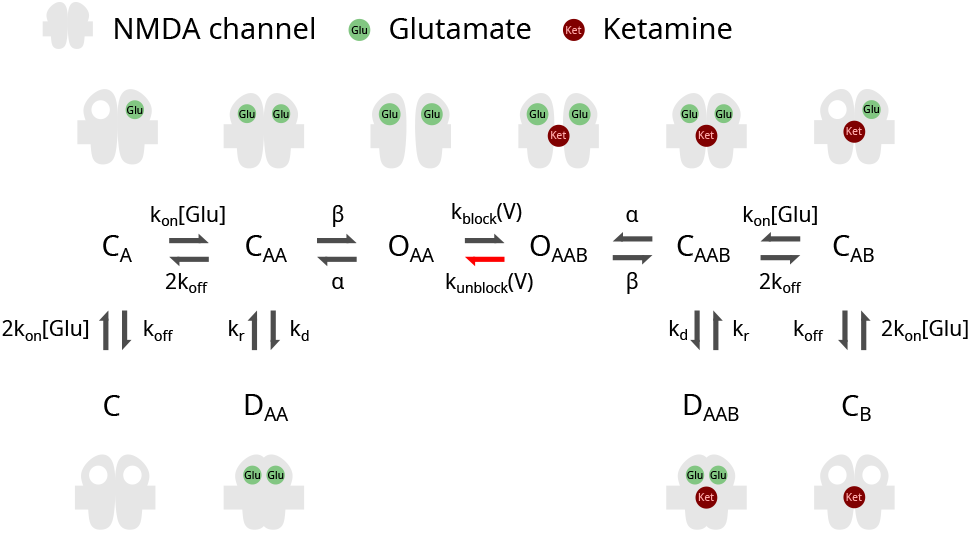
Transitions in the 10-state probabilistic model of NMDA_*R*_ kinetics. Schematic of the 10-state model of NMDA_*R*_ kinetics displaying the variables for the transition rates.

